# The community of Marine Alveolate parasites in the Atlantic inflow to the Arctic Ocean is structured by season, depth and water mass

**DOI:** 10.1101/2024.03.01.582906

**Authors:** Elianne Egge, Daniel Vaulot, Aud Larsen, Bente Edvardsen

## Abstract

The marine alveolates (MALVs) are a highly diverse group of parasitic dinoflagellates, which may regulate populations of a wide range of hosts, including other dinoflagellates, copepods and fish eggs. Knowledge on their distribution and ecological role is still limited, as they are difficult to study with morphological methods. In this work, we describe the taxonomic composition, seasonal- and depth distribution of MALVs in the Arctic Ocean west and north of Svalbard, based on metabarcoding data from five cruises. We recovered amplicon sequence variants (ASVs) representing all major groups previously described from environmental sequencing studies (Dino-Groups I-V), with Dino-Groups I and II being the most diverse. The community was structured by season, depth, and water mass. In the epipelagic zone, the taxonomic composition varied strongly by season, however there was also a difference between Arctic and Atlantic water masses in winter. The spring and summer epipelagic communities were characterized by a few dominating ASVs, which were present in low proportions during winter and in mesopelagic summer samples, suggesting that they proliferate under certain conditions, e.g., when specific hosts are abundant. The mesopelagic samples were more similar across sampling months, and may harbor parasites of deep-dwelling organisms, little affected by season.

## Introduction

Parasitism is one of the most successful life strategies on earth, as evidenced by its independent evolution in most branches in the tree of life (Poulin and Morand 2000). In the marine environment, parasitism by unicellular eukaryotes (protists) may regulate population sizes of both primary producers and consumers, and thus impact the food web structure and the carbon cycle (Berdjeb et al. 2018; Skovgaard 2014). Consequently, new knowledge on the diversity and host preferences of marine protistan parasites ultimately has the potential to inform and improve models of trophic transfer and biogeochemical cycling (Anderson et al. 2024).

The enigmatic Marine Alveolates (commonly referred to as MALVs) is a polyphyletic assemblage of parasitic dinoflagellates which are considered to be the dominating group of parasites in marine microbial food webs (Anderson et al. 2024; Bjorbækmo et al. 2019; Holt et al. 2023). In environmental sequencing studies, MALVs are generally reported to have high abundance and richness in marine pelagic environments (e.g. Guillou et al. 2008; Koid et al. 2012; Massana et al. 2011; Vargas et al. 2015), including polar waters (Clarke et al. 2019; Cleary and Durbin 2016; López-García et al. 2001; Lovejoy et al. 2006). The existence of high genetic diversity within this group is mostly inferred from environmental sequencing, where a high number of unique rRNA SSU gene sequences are phylogenetically placed next to parasitic dinoflagellate species (Groisillier et al. 2006; Guillou et al. 2008; Holt et al. 2023). Such parasitic dinoflagellates were first described in the 1920s (Chatton 1920) and later in the 1960s (e.g. Cachon 1964). Among the described genera are *Amoebophrya*, which infects other dinoflagellates (Coats and Park 2002), *Syndinium*, known to cause mortality in several copepod species (Kimmerer and McKinnon 1990; Skovgaard et al. 2005), *Hematodinium*, which includes at least three species infecting a wide range of crustaceans, impacting fisheries and aquaculture (e.g., Davies et al. 2022), and *Ichthyodinium* which infects fish eggs. Altogether, World registry of marine species (WoRMs) and AlgaeBase list 43 morphologically described species, but not all of them have a known reference sequence. Due to low visibility within the host and the small size of the free-living dinospores, culture-independent methods including molecular tools are often required to study the MALVs. Such techniques have identified host-specific infections in high-biomass blooms of dinoflagel-lates, (Chambouvet et al. 2008). MALV sequences have also been obtained from single-cell isolates of diatoms, concurrently with observations of putative parasitic cells within diatom cells, which suggests that diatoms also may be infected (Sassenhagen et al. 2020). In addition to parasitism, symbiotic relationships between this group and other protists have been proposed (Worden et al. 2015).

The life-cycle of the described parasitic MALV species is typically an alternation between a bi-flagellated infectious spore stage and an intracellular stage, during which the infecting spores divide and grow to a multinucleated structure which can fill the whole host cell (Cachon 1964). For *Amoebophrya* the intracellular stage is called the trophont, and the maturation of the trophont lasts 2-3 days. Upon release from the host, the trophont elongates to form a swimming structure called the vermiform, which may be larger than 50 *µ*m. Within a few hours, the vermiform disintegrates into the free-living dinospores (Chambouvet et al. 2008) which are usually 1-12 *µ*m. Species of *Syndinium* have a similar cycle, but the vermiform stage has not been reported, instead the spores escape as free-swimming zoospores upon release from the host (Skovgaard et al. 2005). For some *Syndinium* species the spores may be up to 20 *µ*m in diameter. Infection by MALVs almost always kills the host, except for some larger crustaceans (Davies et al. 2022).

The MALV group was long thought to be a monophyletic sister group to the dinokaryotes (i.e., dinoflagellates with permanently condensed chromosomes, Orr et al. 2012), and is often collectively referred to as “Syndiniales” (Holt et al. 2023). Based on the rRNA SSU gene, the MALV group is split into two main groups, known in the literature as MALV I and II or Dino-Group (DG) I and II (Groisillier et al. 2006; Guillou et al. 2008). Guillou et al. (2008) also established the smaller clades DG-III - V. Since then, the MALV has been shown to be polyphyletic, with DG-II and DG-IV forming a sister clade to the dinokaryotes and DG-I placed basal to these clades in the dinoflagellate phylogeny (Strassert et al. 2018). Recently, phylogenomics based on single-cell transcriptomics has shown that DG-I and DG-II/IV evolved independently from two distinct free-living ancestors (Holt et al. 2023). The described genera in DG-I are *Ichthyodinium*, *Euduboscquella* and *Duboscquella*, the latter two genera have been isolated from ciliate and other dinoflagellate hosts. *Amoebophrya* and *Hematodinium*, which are both characterised by the lack of any trace of a plastid are placed in DG-II/IV, along with *Syndinium* (*Amoebophrya* in DG-II and *Hematodinium* and *Syndinium* both in DG-IV). Holt et al. (2023) suggest retaining the name “Syndiniales” for the group DG-II/IV, and using the order “Ichthyodinida” for DG-I. No described species has been assigned to DG-III and DG-V, and it is not yet clear whether these groups also belong to “Ichthyodinida”. Based on SSU rDNA phylogenies, DG-I and II/IV are further divided into 8 and 57 subclades, respectively, but the relationship between these clades and described genera is sometimes unclear, e.g., *Amoebophrya* sequences are placed in several subclades (Guillou et al. 2008). In the taxonomy of the Protist Ribosomal Reference database (PR^2^, Guillou et al. 2013; Vaulot 2022), the different Dino-Groups are assigned at the “order” level, and the subclades are assigned at the “family” level. In the literature these subclades are numbered and referred to as e.g., Dino-Group I Clade 1, and we will hereafter refer to them as “clades”. DG-II/IV is generally considered the most diverse since it has the highest number of clades, and also the highest fraction of environmental sequences assigned to it (Table 1, Groisillier et al. 2006; Guillou et al. 2008). Communities of microbial parasites and MALV in particular in a given layer of the water column may be shaped by biological factors such as access to hosts, and physical factors such as mixing or stratification of the water column, and sinking when attached to hosts or particles (Anderson et al. 2024). Early meta-studies of environmental sequencing datasets suggested that MALV clade composition differs between the photic and aphotic zone (Guillou et al. 2008). A recent metabarcoding study of MALV composition in the water column in the Sargasso sea found depth-structuring which was repeated over several years, and in this temperate region the MALV community was more strongly structured by depth than season (Anderson et al. 2024). However, the seasonal variation of the pelagic MALV community composition is not well studied (Anderson and Harvey 2020), especially in polar regions and at mesopelagic depths, where sampling during the winter is challenging due to darkness and rough weather. In the Arctic, seasonality in primary production, and thus in protist assemblages, is strong in the epipelagic waters due to the extreme variation in light regime (e.g. Marquardt et al. 2016). Furthermore, during the spring bloom, zooplankton such as copepods migrate to the surface to feed on the phytoplankton bloom (Hobbs et al. 2020). Thus, there is high seasonal variability in the assemblage of potential MALV hosts in this environment. The mesopelagic zone in the Arctic has been much less studied, and little knowledge is available on parasite-host interactions in the deep ocean. The mesopelagic zone is of particular interest regarding carbon sequestration, as it is an important processing zone for sedimenting euphotic production (Terrado et al. 2009). Microbially mediated processes in these deep waters are considered to be crucial for organic matter remineralization, and thus impact the oceanic carbon pump (Nagata et al. 2010). To understand the impact of parasitism by MALV on the marine food web and carbon cycling in the Arctic, knowledge of the dynamics and distribution with time and depth is necessary.

**Table 1.**
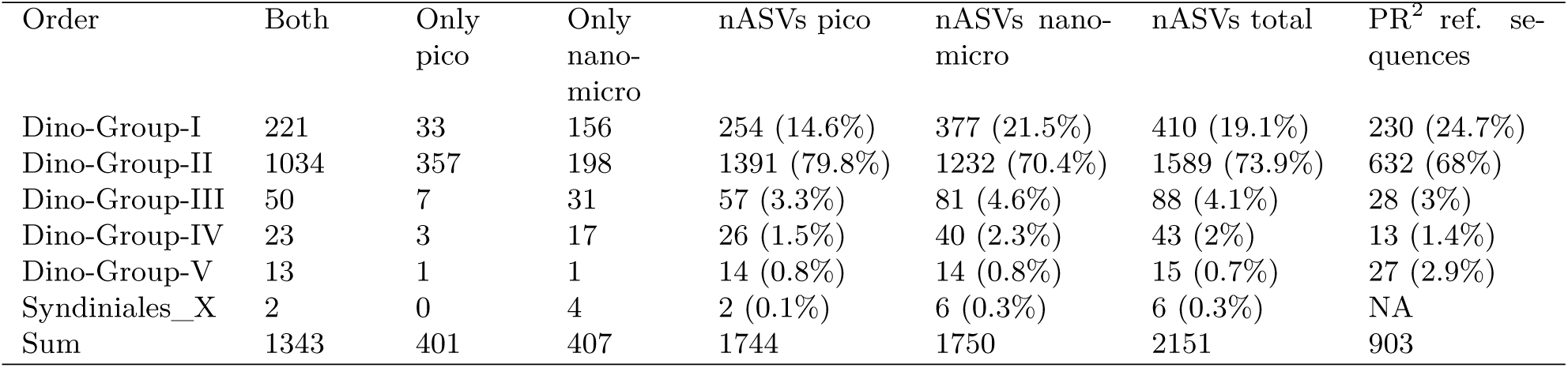
Distribution of ASVs between the Dino-Groups, in both size fractions. “Both” = Number of ASVs found in both size fractions. “Only pico” = Number of ASVs only found in the pico fraction (0.4-3 *µ*m). “Only nano-micro” = Number of ASVs only found in the nano-micro fraction (3-200 *µ*m). The taxonomic distribution of the PR^2^ reference sequences is given for comparison. nASVs= number of ASVs. Syndiniales_X are unclassified MALV sequences falling outside the known Dino-Groups.

| Order | Both | Only<br>pico | Only<br>nano-micro | nASVs pico | nASVs nano-micro | nASVs total | PR<sup>2</sup> ref. se-<br>quences |
| --- | --- | --- | --- | --- | --- | --- | --- |
| Dino-Group-I | 221 | 33 | 156 | 254 (14.6%) | 377 (21.5%) | 410 (19.1%) | 230 (24.7%) |
| Dino-Group-II | 1034 | 357 | 198 | 1391 (79.8%) | 1232 (70.4%) | 1589 (73.9%) | 632 (68%) |
| Dino-Group-III | 50 | 7 | 31 | 57 (3.3%) | 81 (4.6%) | 88 (4.1%) | 28 (3%) |
| Dino-Group-IV | 23 | 3 | 17 | 26 (1.5%) | 40 (2.3%) | 43 (2%) | 13 (1.4%) |
| Dino-Group-V | 13 | 1 | 1 | 14 (0.8%) | 14 (0.8%) | 15 (0.7%) | 27 (2.9%) |
| Syndiniales_X | 2 | 0 | 4 | 2 (0.1%) | 6 (0.3%) | 6 (0.3%) | NA |
| Sum | 1343 | 401 | 407 | 1744 | 1750 | 2151 | 903 |

The present study was conducted in the northern Svalbard region of the Arctic Ocean which was sampled in 2014 during five cruises representing the full seasonal cycle, at three to four depths from the surface down to 1000 m. Seawater samples were size-fractionated into two to four size fractions between 0.4-200 *µ*m (including pico-, nano- and micro-plankton). The taxonomic composition of the protist community was determined from metabarcodes (Amplicon Sequence Variants or ASVs) of the 18S rRNA gene V4 region. The full dataset is presented in Egge et al. (2021). In the present paper, we focus on the MALV community to shed light on the processes that drive its diversity and distribution at epi- and mesopelagic depths in the Arctic Ocean. Our analyses reveal different seasonal distributions in the epipelagic and mesopelagic zones, and different seasonal changes in the pico- and nano-micro fractions. During winter and early spring, we also observed differences in community composition in the epipelagic zone between stations influenced by water flowing south from the Arctic basin, and stations dominated by deep convection of Atlantic water. In spring (May), the photic zone was dominated by a few clades and ASVs which were almost absent in winter and below the photic zone, whereas the aphotic zone community in spring and summer resembled the winter community. We found evidence for a unique MALV community at 1000 m depth, indicated by a high number of ASVs not recovered from other samples. In terms of clade composition, our data are similar to studies from lower latitudes, with high abundance of DG-I-Clade 1 and DG-I-Clade 5 in the photic zone under sunlit conditions, and DG-II-Clade 6 and 7 at mesopelagic depths, suggesting a wide distribution and high degree of adaptation within these clades. Finally, we identified ASVs which are abundant in the surface samples during the spring bloom and which could potentially represent parasites of key species in the Arctic pelagic food web.

## Materials and Methods

A detailed description of sampling, molecular lab work and bioinformatic processing can be found in the data paper from the MicroPolar project (Egge et al. 2021). The methods are briefly repeated here. Data from Egge et al. 2021, including sampling dates, station coordinates and sampled depths, can be found at https://doi.org/10.17882/79823.

### Sampling

Water samples were collected during five cruises west and north of the Svalbard archipelago in the Atlantic and Arctic Oceans in 2014 as part of the MicroPolar project (Paulsen et al. 2016; Sandaa et al. 2018; Wilson et al. 2017). During January (06.01–15.01), March (05.03–10.03), and August (07.08–18.08) cruises, samples were collected from the southern branch of the Western Svalbard Current, which transports water into the Arctic Ocean. During May (15.05–02.06), August (07.08–18.08) and November (03.11–10.11) transects across the core of the Atlantic water inflow were made between 79 *^◦^*N and 79.4 *^◦^*N (Figure 1 A). The sampling area and locations were largely determined by the sea ice cover. During each cruise, 3-6 stations were sampled along a transect at four depths: in the epipelagic zone at 1 m and at a depth between 15-25 m (for the spring and summer samples, this is where the deep chlorophyll maximum, DCM, was found), and in the mesopelagic zone at two depths, in general 500 m and 1000 m, or 10 m above the bottom at shallower stations.

**Figure 1.**
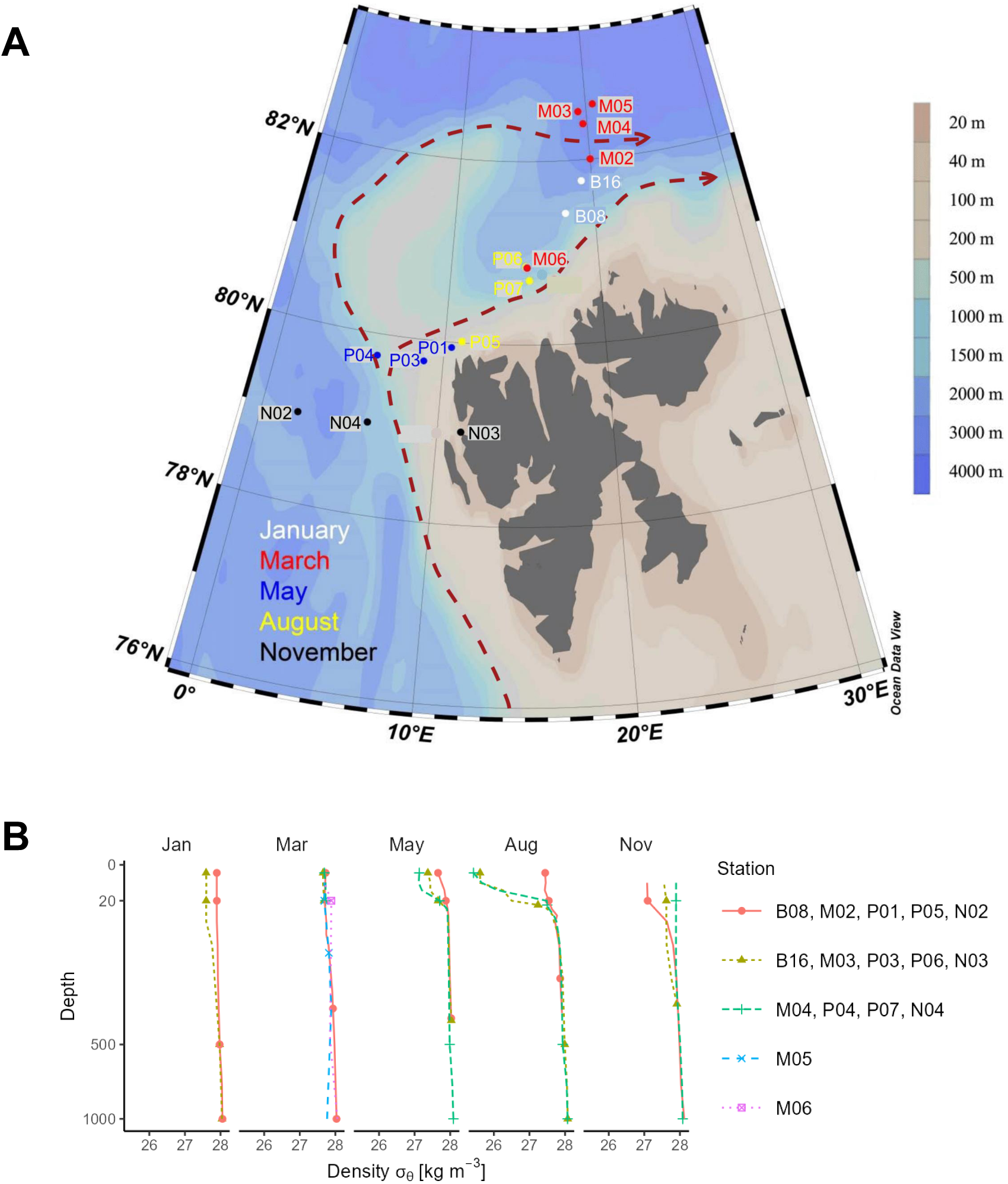
Map of the sampling area and water column density profiles at each sampling. A) Map, sampling stations are indicated with dots, colored according to sampling month, and labelled with names. The arrows show the trajectory of the West Spitsbergen Current, which splits into two branches near the Yermak Plateau. Bottom depth is indicated by the color scale. B) Density profiles of the water column at each station.

### Sampling preparation for DNA extraction

Fifty litres of seawater were sampled from each station and depth. During the January and March cruises, the samples were pre-filtered through a 180 *µ*m mesh, and size fractionated into the 0.4-3 *µ*m (picoplankton) and 3-180 *µ*m (nano- and micro-plankton) fractions by peristaltic pumping (Masterflex 07523-80, Cole Parmer, IL, USA), through serially connected 3 *µ*m and 0.4 *µ*m polycarbonate filters (142 mm diameter, Millipore), mounted in stainless-steel tripods (Millipore, Billerica, MA, USA). The filters were placed in cryovials with AP1 buffer (DNeasy Plant mini kit, Qiagen, Hilden, Germany), flash frozen in liquid N_2_ and kept at -80 *^◦^*C until DNA extraction. During the May, August and November cruises, the water was sequentially poured through 200, 50, and 10 *µ*m nylon mesh, the material on each nylon mesh was collected with sterile filtered seawater into a 50 mL Falcon tube, and collected by filtration on a polycarbonate filter (10 *µ*m pore size, 47 mm diameter, Millipore, USA). The plankton smaller than 10 *µ*m passing through the nylon mesh system was fractionated into the 3-10 *µ*m (small nanoplankton) and 0.4-3 *µ*m size fractions by serial filtration through 142 mm diameter polycarbonate filters. The filters were preserved as described above.

### DNA extraction, PCR and sequencing

DNA was extracted with the DNeasy Plant mini kit (Qiagen, Hilden, Germany), according to the manufacturer’s protocol, except that frozen samples were incubated at 95 *^◦^*C for 15 min, then shaken in a bead-beater 2 x 45-60 seconds to rupture cell coverings. The V4 region of the 18S rRNA gene was amplified with primers TAReuk454FWD1 (5’-CCAGCASCYGCGGTAATTCC-3’) and V4 18S Next.Rev (5’-ACTTTCGTTCTTGATYRATGA-3’) (Piredda et al. 2017). The amplification primers were tagged with internal barcodes to allow sample multiplexing. PCR reactions were performed with KAPA HiFi HotStart ReadyMix 2x (KAPA Biosystems, Wilmington, MA, USA), in triplicate for each sample, and pooled prior to purification and quantification. The PCR products were purified with AMPure XP beads (Beckman Coulter, Brea, USA), quantified with NanoDrop and subsequently pooled in equal concentrations. The pools were sent to library preparation at the Norwegian Sequencing Centre (NSC, Oslo, Norway) and GATC GmbH (Konstanz, Germany) with the KAPA library amplification kit (KAPA Biosystems, Wilmington MA, USA). Due to delivery problems with the Illumina MiSeq chemistry in the spring of 2015, the sequencing was done with a modified HiSeq protocol on two HiSeq runs at GATC. After initial analysis of the HiSeq data, samples with low number of reads were re-sequenced with Illumina MiSeq v.3 (300 bp, paired-end) at the NSC.

### Sequence processing

Reads were processed into Amplicon Sequence Variants (ASVs) and chimeric ASVs were removed using the R library dada2, v1.16. (Callahan et al. 2016), as described in detail in (Egge et al. 2021). The ASVs were taxonomically assigned with function *assignTaxonomy*, the dada2 implementation of the naive Bayesian classifier method (Wang et al. 2007), against the PR^2^ database (Guillou et al. 2013; Vaulot 2022), version 4.12.0 (https://github.com/pr2database/pr2database/releases/tag/v4.12. 0). Amplicon sequence variants with less than 90% bootstrap value at class level and/or which comprised less than 10 reads in total were removed. In PR^2^, sequences which are assigned to a taxonomic group at a given level, but which cannot be assigned to lower taxonomic levels are denoted with “_X“, e.g., DG-I_X, and we will follow this notation. In the current version of the PR^2^, “Syndiniales” is still used as a synonym for the entire MALV group, and thus reference sequences in PR^2^ which could not be assigned to any of the Dino Groups are assigned to “Syndiniales_X”. We therefore use this group name in the results.

### Statistical analyses

All statistical analyses were performed in R (version 3.3.1, http://r-project.org), with the packages phyloseq, v. 1.38.0 (McMurdie and Holmes 2013), microbiome, v. 1.12.0 (Lahti and Shetty 2012-2019) vegan 2.5-7 (Oksanen et al. 2021), and mvabund 4.1.12 (Wang et al. 2022), unless otherwise stated. Figures were created with the r-packages ggplot2 (Wickham 2016) and ragg (Pedersen and Shemanarev 2022).

### Preparation of MALV ASV tables

For the picoplankton size fraction (0.4-3 *µ*m), MALV ASVs were extracted from the 0.4-3 *µ*m samples in the original ASV table (i.e., not rarefied, entries given as read counts). To be able to compare the May, August and November samples between 3 *µ*m and 200 *µ*m with the January and March 3-180 *µ*m samples, we merged these into the size fraction 3-200 *µ*m, in the following referred to as ‘nano-micro‘. To ensure proportionate contribution of reads from each size fraction, the total protist ASV table was subsampled to a number of reads corresponding to the sample with fewest reads, as follows: 88,000 reads from 3–180 *µ*m; 40,000 reads from each of 3–10 *µ*m and 10–50 *µ*m, and 8,000 reads from 50–200 *µ*m. Subsampling to equal read number was performed 100 times, and the average read number per ASV was used, rounded to 0 decimals. Subsampling was done with the function *rrarefy* from the vegan package. The low number of protist reads in the 50–200 *µ*m fraction was due to a high proportion of Metazoan reads in this fraction. MALV ASVs were then extracted from these rarefied samples, and the the fractions 3-10, 10-50 and 50-200 *µ*m were merged into the fraction 3-200 *µ*m by taking the sum of the read number for each ASV.

### Alpha diversity

For the alpha diversity analyses of the MALV community (richness, evenness and Shannon diversity), all samples in the pico- and nano-micro size fractions were subsampled to 5,000 MALV reads, with the *rrarefy*-function, as described above. ASV richness and Shannon diversity were calculated with the functions *specnumber* and *diversity*, respectively. As recommended by Borcard et al. (2018), Shannon diversity is given as the Hill number *N*_1_ (= *exp(H)*, where *H* is Shannon entropy), expressed as an ASV number equivalent. *N*_1_ can then be interpreted as the number of ASVs needed to obtain the observed Shannon entropy, if all ASVs had equal proportion. Evenness is defined as *N*_1_/*N*_0_, where *N*_0_ is the number of observed ASVs.

### Beta diversity

For the beta diversity analyses (i.e., comparisons of community composition), we used the un-rarefied MALV ASV tables, following recommendations by McMurdie and Holmes (2014). Amplicon sequencing data are inherently compositional, i.e., the proportions of the different ASVs are not independent of each other (Gloor et al. 2017). This may lead to spurious correlations between ASVs, and incorrect estimates of similarity between samples (Jackson 1997). According to the recommendations of Gloor et al. (2017), the MALV read counts were therefore transformed to centred log-ratio values (CLR), with the function *transform* from the microbiome package. Prior to calculating the log-ratio, this function transforms zeroes by adding the smallest ASV-proportion in the whole dataset, divided by two (Lahti and Shetty 2012-2019). CLR-transformations are made by calculating the geometric mean of the read counts of all the ASVs within a sample, dividing the read count of each ASV on this geometric mean and log-transforming (Nearing et al. 2022). Thus, similar to the original proportion, the CLR value indicates how dominating an ASV is in a particular sample. In addition, CLR values for a given ASV can be compared between samples (Gloor et al. 2017). Similarity in CLR-transformed ASV-composition between samples was estimated by the Aitchison distance. This distance is defined as the euclidean distance between CLR-transformed samples, and was calculated with the function *vegdist* in package vegan. Principal component analysis was then performed with the function *prcomp*. The samples were clustered according to Aitchison distance with the function *hclust* with complete linkage clustering. Clusters of interest were then delineated after visual inspection of the dendrogram and PCA plots, to identify clusters that corresponded to certain environmental conditions, e.g., combinations of season and depth. Clade composition was visualised as bar charts and the CLR of dominating ASVs were visualised as heatmaps. Plots of shared ASVs between sample clusters were created with the package UpSetR v. 1.4.0 (Conway et al. 2017). To assess which ASVs were significantly differentially distributed between these clusters, we used the function *anova.manylm* from the mvabund package. This function fits multivariate linear models to the CLR-transformed abundance table and tests whether the ASVs have significantly different CLR-values between groups of samples (in this case clusters). Homogeneity of variances was checked by plotting residuals against the fitted values. The function *anova.manylm* was performed 10 times, and ASVs which had p-value < 0.05 in 5 or more of the trials were considered to have significant differential abundance between the clusters.

### Phylogenetic analyses

To assess phylogenetic relationships below clade level between the most abundant ASVs, the most abundant ASVs assigned to the abundant clades DG-I-Clade-1, DG-I-Clade-5, DG-II-Clade-6 and DG-II-Clade-7 were aligned with MAFFT v. 7, with the G-INS-1 option (mafft.cbrc.jp; (Katoh et al. 2019)) and trimmed with trimAl (Capella-Gutiérrez et al. n.d.). The phylogenetic tree was created with RAxML based on 340 sites and 100 times bootstrap resampling. The tree was imported into R using functions from the treeio package (Wang et al. 2020), and visualised with functions from the package phyloseq.

### Biogeographic distribution of ASVs and clades

The biogeographic distribution of particular ASVs and clades of interest was investigated using the metaPR^2^ online shiny application (Vaulot et al. 2022, ; https://shiny.metapr2.org, version 1.0). For each ASV, the sequence was used as query in a BLAST-like search, and the biogeographic distribution was taken as the combined distribution of the metaPR^2^ ASVs with 100% identity to the queried ASV. Similarly, clade distribution was assessed by using the taxon search.

## Results and discussion

### High taxonomic diversity of MALV in the Arctic

MALV was the most diverse protistan group in the metabarcoding dataset, constituting 33% of the unique amplicon variants (ASVs). It was the most abundant group in the picoplankton size fraction (0.4-3 *µ*m), where it constituted between 6 and 99% of the reads in each sample (Egge et al. 2021, Figure S1). We recovered in total 2,151 MALV ASVs. Dino-Groups I and II were the most diverse, with 410 and 1,589 ASVs, respectively, corresponding to 19% and 74% of the MALV ASVs (Table 1). The order-level clades Dino-Group DG-III, DG-IV, DG-V and “Syndiniales_X” (PR^2^ sequences assigned to MALVs which cannot be placed in any Dino-Group) had 88, 43, 15 and 6 ASVs, respectively (corresponding to 4, 2, 0.1, 0.7 and 0.3%). This is similar to the taxonomic distribution of the PR^2^ MALV reference sequences (Table 1), but with a higher percentage of DG-II ASVs, and lower percentage of DG-I and DG-V. The number of ASVs was similar in the two size fractions, with 1,744 in the pico fraction (0.4-3 *µ*m) and 1,750 in the nano-micro size fraction (3-200 *µ*m) (Table 1). The pico fraction had higher richness of DG-II and lower of DG-I compared to the nano-micro fraction (80 and 15% of ASVs in this fraction vs. 70 and 22%, respectively). This was mostly due to higher richness of DG-II-Clade 1 in the pico fraction. Higher abundance of DG-II in the pico fraction compared to the larger fractions is also found in the global metaPR^2^ dataset (Vaulot et al. 2022).

At lower taxonomic levels, within DG-I, we detected sequences from all the eight subclades delineated in Groisillier et al. (2006) and Guillou et al. (2008), and in addition 7 ASVs which could not be placed in a clade within DG-I. From DG-II we detected sequences assigned to 40 out of the 57 subclades included in PR^2^ (Guillou et al. 2013), and 139 ASVs which could not be placed in a clade. In both fractions, DG-II-Clade-1 had the highest number of ASVs, with in total 305 ASVs (14.2% of the ASVs) (273 and 205 in the pico- and nano-micro fractions, respectively, corresponding to 16 and 12%). DG-I-Clade 5, DG-II-Clade 10-&-11, DG-II_X and DG-II-Clade 7 had between 160 and 120 ASVs each, corresponding to 7-5.5% (Figure 2, Table S1). Aside from DG-II-Clade 1, the family-level taxonomic distribution of ASVs were similar between the two size fractions (R^2^ = 0.9). Interestingly, there was generally no correlation between the number of ASVs assigned to each clade, and the mean read abundance, in any of the size fractions (R^2^ = 0; Figure 2). For example DG-II-Clade 1, which was the most ASV-rich in both size fractions, had less than 4% mean read abundance. Conversely, DG-I Clade 1 and 5 had lower richness, but high mean read abundance.

**Figure 2.**
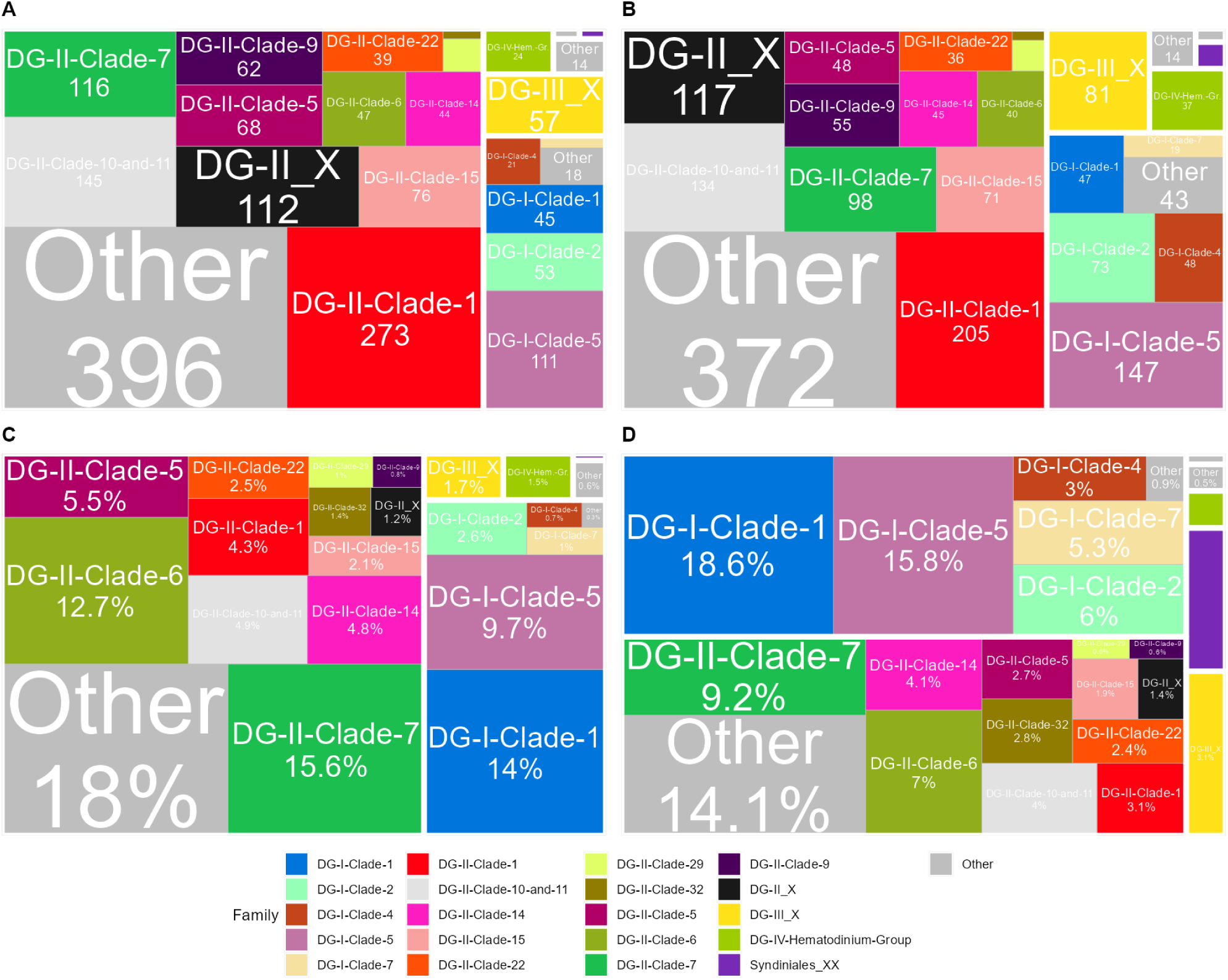
Treemaps of MALV taxonomic composition. Number of ASVs per clade in the pico (0.4-3 *µ*m, A) and nano-micro (3-200 *µ*m, B) size fractions, respectively. Mean percentage of reads per clade in the pico- (C) and nano-micro (D) fractions. Clades represented by less than 10 % of the reads in all samples are grouped into the category “Other”. For information on these clades, see Table S1.

### The MALV community in the Arctic is shaped by season, depth and water mass

The taxonomic composition of the samples was related to a combination of season, depth zone and water mass in both size fractions, as shown by principal component ordination of taxonomic similarity between samples (Figure 3, A and C). Along the first principal component, there was a separation between samples taken in the epipelagic zone in spring and summer, when there was sufficient light for primary production, and samples taken during winter andor in the mesopelagic zone where there was little light. This axis explained 30% and 27% of the variation in ASV composition in the pico- and nano-micro fractions, respectively. Along the second axis, there was a separation within the winter andor mesopelagic samples, where the samples taken at 1000 m were separated from the rest, in both size fractions. Furthermore, the samples from the epipelagic zone in January and March clustered according to the prevailing water mass in the surface. The second axis explained 20 and 24% of the variation in the two fractions, respectively. Complete linkage hierarchical clustering, in combination with visual inspection of the PCA, allowed to delimit six main sample clusters in each of the size fractions. The sample composition in these clusters is described in Table 2. In the pico fraction, 19 ASVs had significantly different CLR-values between the clusters, whereas the nano-micro fraction had 114 (Table S2). Most of these ASVs had higher CLR in the “dark” samples (i.e., samples from the mesopelagic zone, or epipelagic zone in winter; Figure S3).

**Figure 3.**
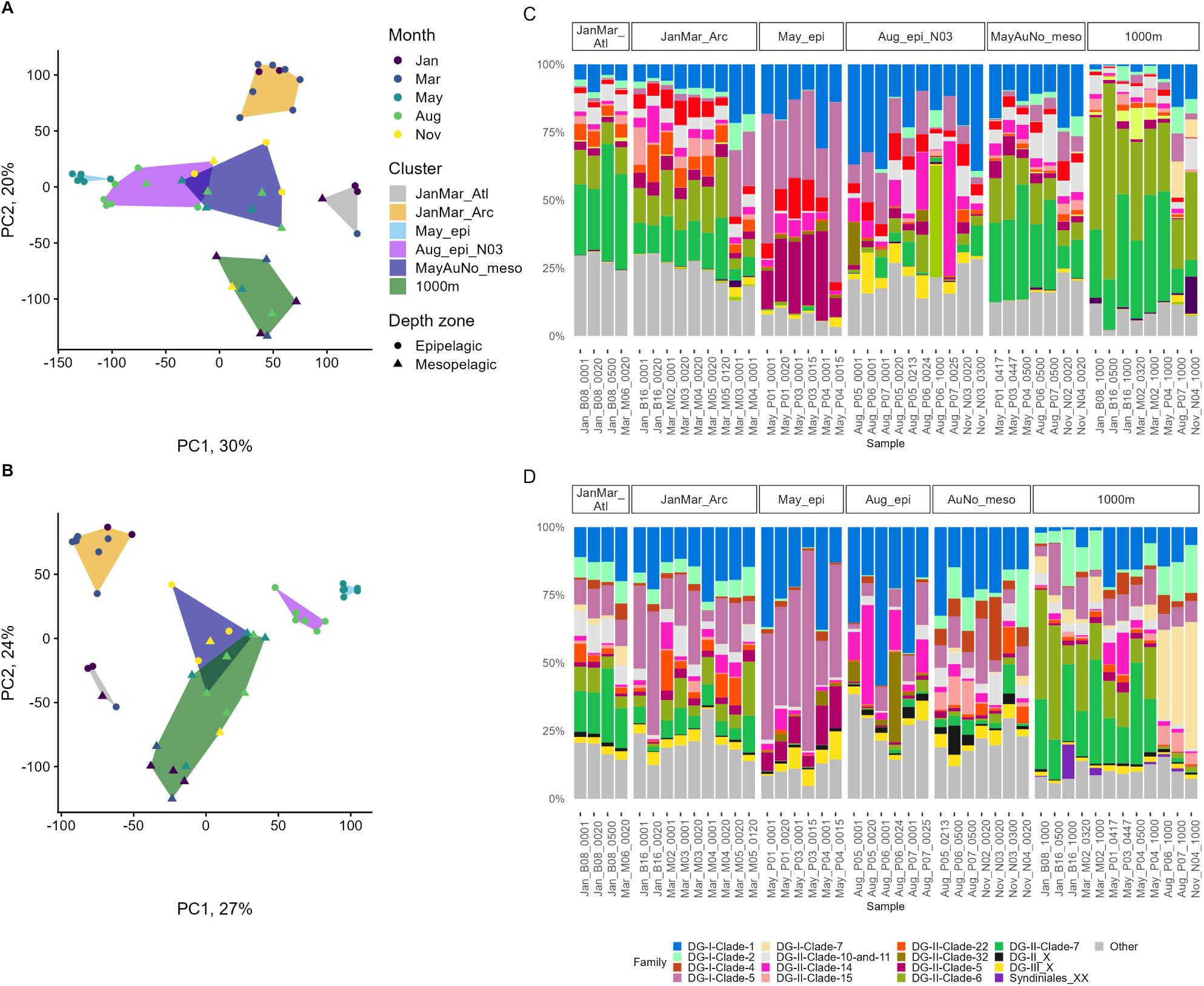
Structuring of MALV community composition. (A) and (C) Principal component analysis of similarity in ASV composition between samples with percent of variation explained by each axis given next to the axis name. (A) pico (0.4-3 *µ*m) fraction, (C) nano-micro (3-200 *µ*m) fraction. (B) and (C) Barplots showing taxonomic composition at the clade level (corresponding to ”family” level in PR^2^). Clades represented by less than 10 % of the reads in all samples are grouped into the category “Other”. The samples are grouped according to the clustering produced by the PCA.

**Table 2.**
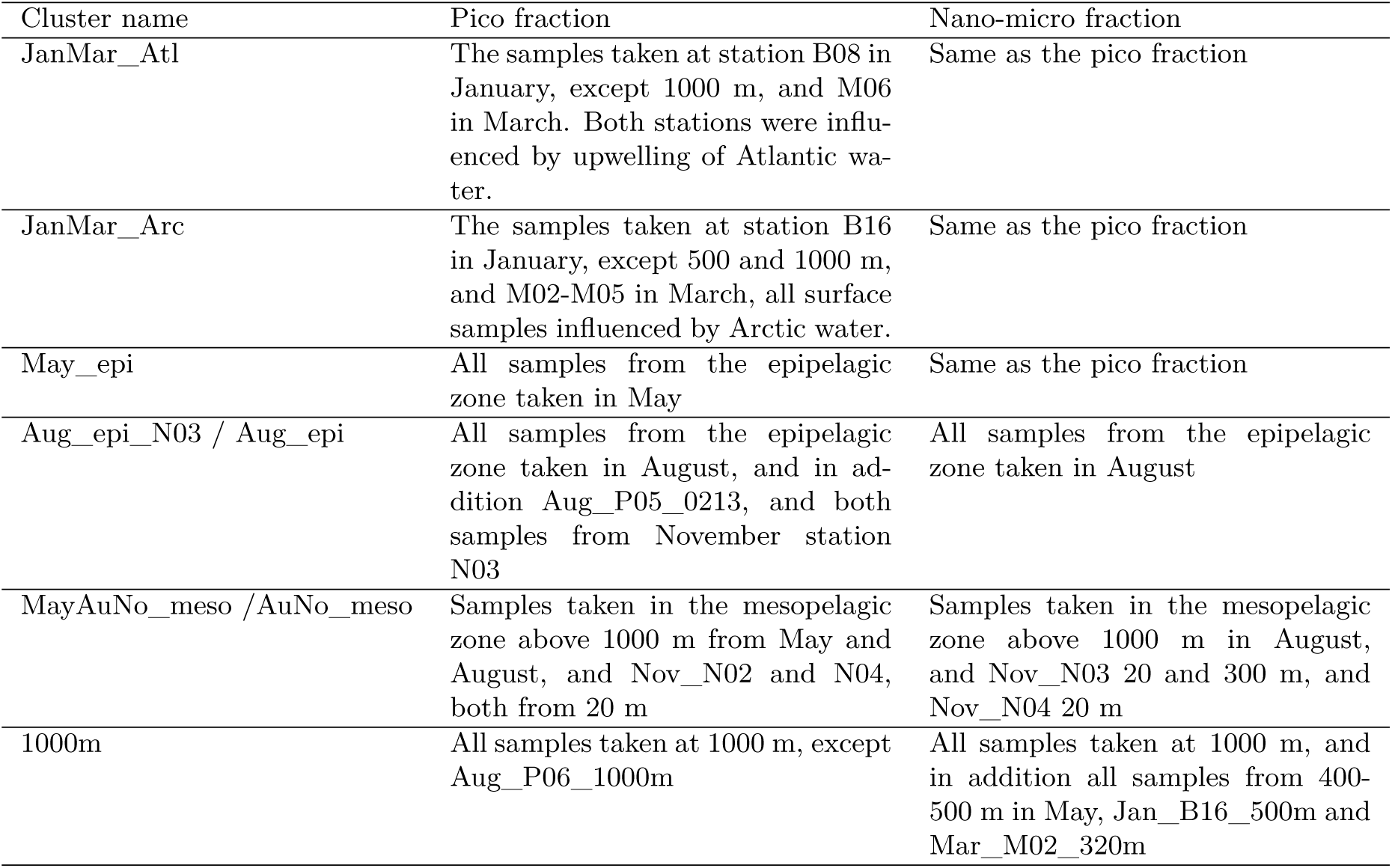
Designation of sample clusters. When 2 names are present in the name column, they correspond to the pico and nano fractions, respectively.

### Evidence for a distinct MALV community at 1000 m

Strikingly, most of the samples taken at 1000 m clustered together and thus had similar ASV composition irrespective of season, in particular in the pico fraction (Figure 3). This suggests that the community of MALV at great depths is relatively stable over time, and may be parasitic of deep-dwelling organisms that are little affected by the strong seasonal variation in the Arctic epipelagic zone. Richness in this cluster was similar to the winter and summer mesopelagic samples above 1000 m, but evenness was lower, in particular in January and March (Figure 4). This cluster also had the highest number of unique ASVs (i.e., only detected in this cluster, Figure S5). A high number of unique MALV ASVs at depths 800 - 1000 m has also been found in oligotrophic, tropical oceans (Anderson et al. 2024). The most abundant clades in this cluster were DG-II Clades 6 and 7, and the most abundant ASVs were also assigned to these clades. Clades 6 and 7 are among the most diverse in DG-II in PR^2^, and Clade 7 has been found to be the most diverse within DG-II in samples from the aphotic zone (Guillou et al. 2008). In metaPR^2^ both DG-II Clade 6 and 7 are detected from tropical to polar latitudes, and are most abundant in mesopelagic samples (Table S3). They are also reported in high proportions in deep samples in Antarctic as well as temperate oceanic regions (Anderson et al. 2024; Cleary and Durbin 2016). In the nano-micro fraction, the August and November 1000 m samples were distinguished by high abundance of DG-I-Clade 7 (mainly in the 3-10 *µ*m fraction), which shows that variation also occurs at these depths. In metaPR^2^, DG-I-Clade 7 is detected from tropical to polar latitudes, generally in low abundance, in the surface as well as at mesopelagic depths. We may thus speculate that this clade infects hosts with vertical migration. ASVs assigned to this clade has been found in high relative abundance in amphipods (Savage et al. 2023). The abundant Arctic amphipod *Apherusa glacialis* is known to perform seasonal vertical migration, and may overwinter in the deep within the Atlantic-water inflow near Svalbard (Drivdal et al. 2021; Kunisch et al. 2020), however, further investigations are needed to establish whether this particular species is susceptible to MALV infection. Other potential hosts at these depths are Arctic copepod species (e.g., *Calanus glacialis* and *C. hyperboreus*) which are known to descend to depths as far as below 800 m for diapause during winter (Kvile et al. 2019, and references therein). *C. hyperboreus* spawns during winter down to 1000 m (Hirche and Niehoff 1996). There is generally little knowledge on the extent of variation in the community of parastic protists at mesopelagic depths in the Arctic. Terrado et al. (2009) found seasonal variation in the protist community at 200 m depth, which the authors suggest was due to advected water masses with a distinct community. However, since we only observed the change in community composition at 1000 m in one size fraction and at geographically separated stations, advection is a less likely explanation in this case.

**Figure 4.**
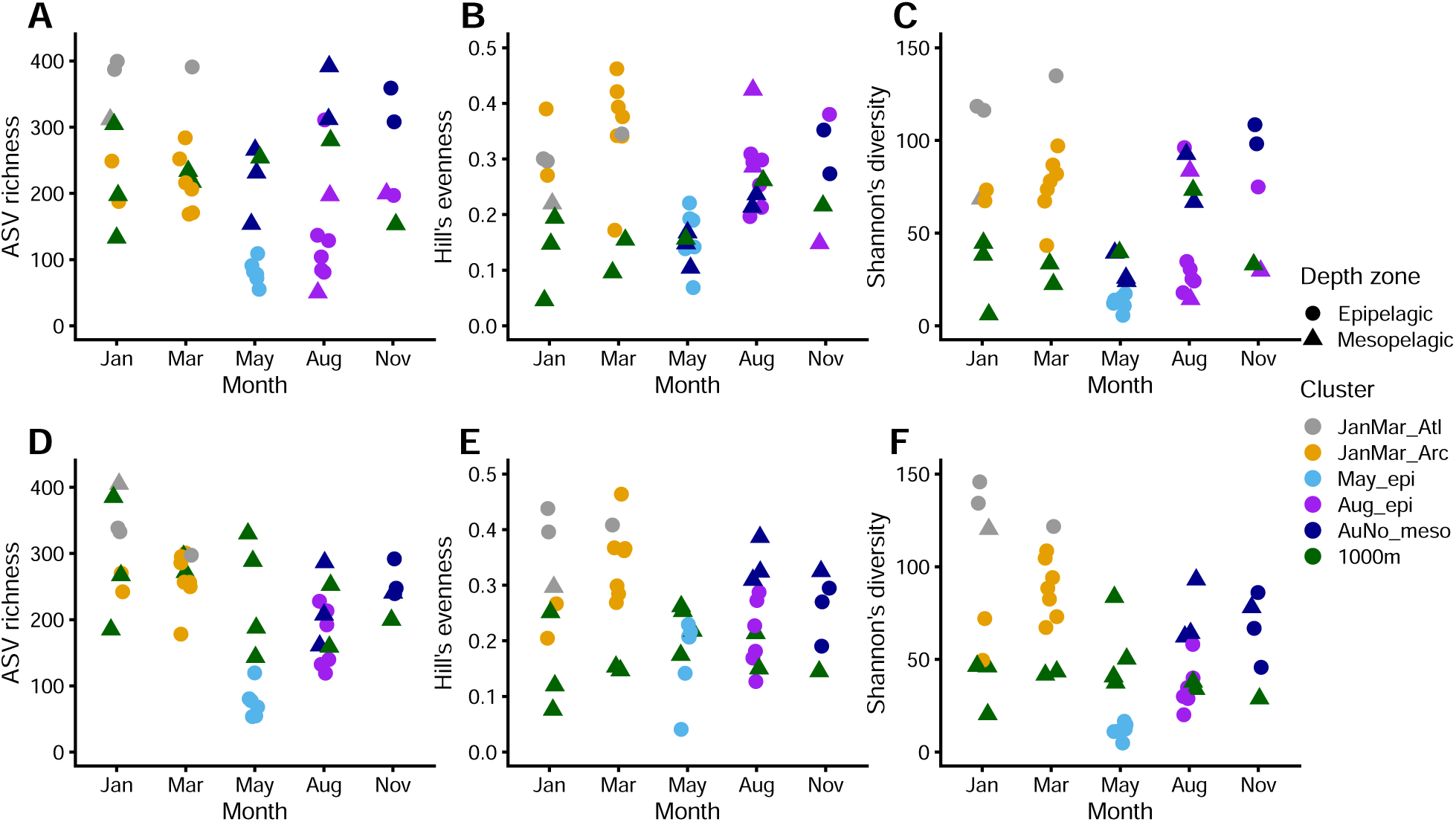
Alpha diversity. (A, D) ASV richness, (B, E) Hill’s evenness, (C, F) Shannon’s diversity. Top row: pico (0.4-3 *µ*m) fraction, bottom row: nano-micro (3-200 *µ*m) fraction.

### Similar communities in samples taken during the winter and below the photic zone in spring and summer

In November and January, polar night conditions (i.e., no daylight) prevailed. In March, day length was 6-8 hours, but the chlorophyll *a* values were < 0.1 *µ*g.L*^−^*^1^, indicating that the spring bloom had not yet started (Figure S2; Randelhoff et al. 2018). The relative abundance of MALV was generally high in both the samples from January, March and November in the epipelagic zone, and in the samples taken in the mesopelagic zone regardless of season, constituting up to 99% of the protist reads in the pico fraction and up to 77% in the nano-micro (Figure S1). High relative abundance of MALV reads in Arctic surface waters during winter has previously been reported from Isfjorden, Svalbard (Marquardt et al. 2016). Furthermore, our Arctic data from mesopelagic depths are consistent with previous studies finding high relative abundance of MALV at great depths in various oceanographic regions (Pernice et al. 2016; Terrado et al. 2009; Xu et al. 2017). At the clade level, the samples taken at mesopelagic depths above 1000 m in summer and winter, and in the winter surface had taxonomic compositions that were a mixture between the surface samples from spring and summer (described below) and the 1000 m samples. Similar to the surface samples from spring and summer, these samples had high proportions of DG-I-Clade 1, and in some samples DG-I-Clade 5, in particular in the nano-micro fraction (Figure 3). Similar to the 1000m cluster, Dino-Group-II was dominating, especially DG-II-Clade 6 and DG-II-Clade 7. DG-II-Clade 7 had the highest number of ASVs with significantly different CLR between clusters in both size fractions, and these ASVs had typically higher CLR-values in the clusters with “dark” samples (Figure S3, discussed below).

So far no representative of the deep-dwelling DG-II-Clade 6 and DG-II-C 7 have been isolated, and their hosts are unknown. Members of DG-II-Clade 7 are speculated to be able to parasitize deep planktonic organisms belonging to Cercozoa and Radiolaria (Anderson et al. 2024; Guillou et al. 2008), which are known to harbour a wide diversity of MALV parasites (Bråte et al. 2012). In the overall protist community, the abundance of Cercozoa and Radiolaria was generally low (Figure S1, Egge et al. 2021). However, within Radiolaria, taxonomic composition indeed differed between the spring and summer epipelagic clusters and the other samples: RAD-A, B and C clades were more abundant in the winter samples and samples from the mesopelagic zone during summer, in particular RAD-B-Group IV and RAD-C. At depths below 200 m, the group Chaunacanthida constituted a high proportion of Radiolarian reads (Egge et al. 2021).

### Difference between Arctic and Atlantic water masses in the surface during winter

The MALV communities in the epipelagic and mesopelagic zones during winter were relatively similar (Figure 3). However, we found geographical differences which could be related to the origin of the water mass at the sampling station. In January and March 2014, there was still open water along the north-west coast of Svalbard. During winter, the water masses on the shelf slope north of Svalbard is dominated by Atlantic water with strong vertical convection, whereas the water masses further off the slope is characterised by a surface layer of fresher Arctic water and some stratification even in the winter (Randelhoff et al. 2018). At the station January B08, which was located closer to the shelf compared to January B16, the MALV communities at 1, 20 and 500 m depth were similar. At B16, which was influenced by Arctic water, as indicated by lower salinity in the surface (Figure 1 B), there was a difference in the community between the epi- and mesopelagic zones. At the March station M06, which was located closer to the shelf, the MALV community in the M06_20m sample (the only sample from this station) was similar to the B08 samples from 1-500 m. The other March stations, which were located further North-East, had a fresher layer down to below c. 150 m, and the samples from this layer were similar to the surface samples from Jan B16. The MALV community in the putative Arctic water masses during winter were characterised by higher relative abundance of DG-II-Clade 1 and DG-II-Clade 22 compared to the winter community of Atlantic origin. Also the overall protist community differed between these water masses, with higher abundance of Picozoa in the samples taken further off the shelf.

### Community composition in the epipelagic zone - change between May and August

The May and August samples were collected at a time of continuous daylight in this region. The May_epi and Aug_epi sample clusters were characterised by relatively low ASV richness and low to intermediate evenness, resulting in low Shannon diversity (Figure 4). The clades with the highest mean relative abundance, in both size fractions, were DG-I-Clade-1 (22% and 26% of the reads in the pico- and nano-micro fractions, respectively), DG-I-Clade-5 (20 and 19%), DG-II-Clade-14 (8.3 and 5.7%), and DG-II-Clade-1 (6.3 and 4.8%). In addition, DG-II-Clade-5 was abundant in the pico fraction (9.4%), and DG-III_X in the nano-micro (4.2%) (Figure 3, Table 3). This cluster was also characterised by low proportions of DG-II Clades 6 and 7 compared to the clusters with samples collected during winter and/or in the mesopelagic zone. DG-I-Clade 5 has been found to be one of the most abundant DG-I clades in the euphotic zone, and DG-II-Clade 5 is also mainly found in the same layer (Guillou et al. 2008). Members of DG-I have been associated with diatoms (Sassenhagen et al. 2020). Inspection of the metabarcode database metaPR^2^ revealed that DG-I-Clade 5 occurs in high abundance from tropical to polar latitudes, and is proportionally more abundant in euphotic zone samples (Table S3). It has also been found in high relative abundance in the surface in oligotrophic, tropical oceans (Anderson et al. 2024). DG-II-Clade 5 is detected in the metaPR^2^ database from tropical to polar latitudes, in low relative abundance, but relative abundance seems to increase with latitude (Table S3). DG-I-Clade 1 was also abundant in this cluster, in addition to in the MayAuNo_meso cluster (discussed below). DG-I-Clade 1 is the most diverse clade within Dino-Group-I, and has been retrieved from a wide range of marine habitats (Groisillier et al. 2006). It was found to be particularly abundant in spring and summer in a metabarcoding study of a temperate coastal environment (Yellow Sea, China, Fu et al. 2020). In metaPR^2^, DG-I-Clade 1 has a wide latitudinal distribution, and is most abundant in euphotic zone samples (Table S3). In the August epipelagic samples, DG-II-Clade 14 was abundant. This clade has been found both in oceanic surface and deep waters (Groisillier et al. 2006). In metaPR^2^ it was detected from tropical to polar latitudes, generally in low abundance (Table S3). Members of this clade have been demonstrated to infect *Heterocapsa triquetra* (Chambouvet et al. 2008), a thecate dinoflagellate which may be abundant in Arctic waters (e.g. Seuthe et al. 2011). In the metabarcoding data, the genus *Heterocapsa* was relatively abundant in the fractions smaller than 10 *µ*m all year (Egge et al. 2021). Stratification began to develop in May and was strong in August at the two northernmost stations (P06 and P07). Parasites, either in the form of infected hosts or spores, may aggregate on particles and sink out of the photic zone (Anderson et al. 2024), which would then result in similar compositions of the environmental DNA (eDNA) pools in these zones. However, in May and August the communities in the epi- and mesopelagic zone were very different. Depth-structuring of marine protist communities in general, and MALV communities in particular, has also been found at lower latitudes (Anderson et al. 2024; Ollison et al. 2021). Our results thus support the hypothesis that the MALV community in the mesopelagic zone, as revealed by analysis of eDNA, is shaped by the environment and availability of hosts at these depths, rather than passive sinking transport of surface-dwelling taxa (Anderson et al. 2024).

**Table 3.**
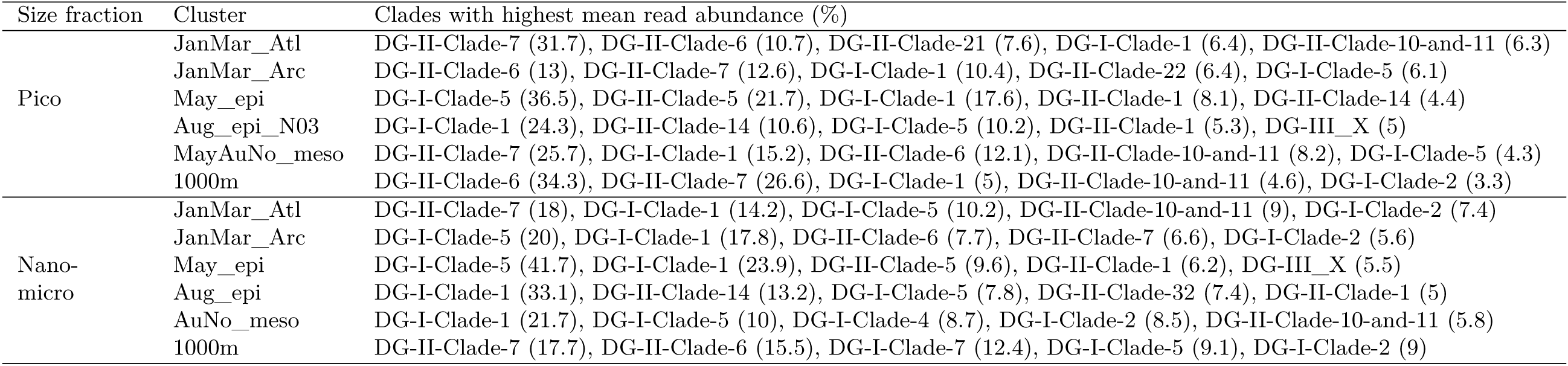
Table of the most abundant clades within each sample cluster shown in Figure 4.

| Size fraction | Cluster | Clades with highest mean read abundance (%) |
| --- | --- | --- |
| Pico | JanMar_Atl | DG-II-Clade-7 (31.7), DG-II-Clade-6 (10.7), DG-II-Clade-21 (7.6), DG-I-Clade-1 (6.4), DG-II-Clade-10-and-11 (6.3) |
|  | JanMar_Arc | DG-II-Clade-6 (13), DG-II-Clade-7 (12.6), DG-I-Clade-1 (10.4), DG-II-Clade-22 (6.4), DG-I-Clade-5 (6.1) |
|  | May_epi | DG-I-Clade-5 (36.5), DG-II-Clade-5 (21.7), DG-I-Clade-1 (17.6), DG-II-Clade-1 (8.1), DG-II-Clade-14 (4.4) |
|  | Aug_epi_N03 | DG-I-Clade-1 (24.3), DG-II-Clade-14 (10.6), DG-I-Clade-5 (10.2), DG-II-Clade-1 (5.3), DG-III_X (5) |
|  | MayAuNo_meso | DG-II-Clade-7 (25.7), DG-I-Clade-1 (15.2), DG-II-Clade-6 (12.1), DG-II-Clade-10-and-11 (8.2), DG-I-Clade-5 (4.3) |
|  | 1000m | DG-II-Clade-6 (34.3), DG-II-Clade-7 (26.6), DG-I-Clade-1 (5), DG-II-Clade-10-and-11 (4.6), DG-I-Clade-2 (3.3) |
| Nano-micro | JanMar_Atl | DG-II-Clade-7 (18), DG-I-Clade-1 (14.2), DG-I-Clade-5 (10.2), DG-II-Clade-10-and-11 (9), DG-I-Clade-2 (7.4) |
|  | JanMar_Arc | DG-I-Clade-5 (20), DG-I-Clade-1 (17.8), DG-II-Clade-6 (7.7), DG-II-Clade-7 (6.6), DG-I-Clade-2 (5.6) |
|  | May_epi | DG-I-Clade-5 (41.7), DG-I-Clade-1 (23.9), DG-II-Clade-5 (9.6), DG-II-Clade-1 (6.2), DG-III_X (5.5) |
|  | Aug_epi | DG-I-Clade-1 (33.1), DG-II-Clade-14 (13.2), DG-I-Clade-5 (7.8), DG-II-Clade-32 (7.4), DG-II-Clade-1 (5) |
|  | AuNo_meso | DG-I-Clade-1 (21.7), DG-I-Clade-5 (10), DG-I-Clade-4 (8.7), DG-I-Clade-2 (8.5), DG-II-Clade-10-and-11 (5.8) |
|  | 1000m | DG-II-Clade-7 (17.7), DG-II-Clade-6 (15.5), DG-I-Clade-7 (12.4), DG-I-Clade-5 (9.1), DG-I-Clade-2 (9) |

There was a difference in the composition of ASVs as well as of clades between the surface samples from May and August. The May_epi cluster was characterised by high abundance of DG-I-Clade 5 and DG-II-Clade 5 (Figure 3), due to the very abundant ASVs ASV_3_DG-I-C5 and ASV_7_DG-II-C5, which constituted up to 70% and 26% of the reads, respectively, in these samples (Figure 5). These ASVsOTUs have previously only been detected in surface samples (Table S4). In the nano-micro fraction, ASV_15_DG-II-C1 was also abundant in the May samples. These ASVs had generally lower abundance in the August samples. Other ASVs showed the opposite pattern, with lower abundance in May than in August, such as ASV_13_DG-I-C5 (pico), and ASV_25_DG-I-C1 and ASV_17_DG-II-C14 (nano-micro). This indicates that the MALV community also follows the succession of the phytoplankton community from spring to summer, possibly as a response to a change in the host community. Strong seasonality of MALV taxonomic composition in sunlit waters has previously been observed in a coastal pond (Sehein et al. 2022) and in subtropical coastal waters, where summer samples were distinct from those of colder months (Anderson and Harvey 2020). The low richness and evenness in May could reflect a host community dominated by few taxa. These samples had high levels of Chl *a* (up to 15 *µ*g L*^−^*^1^; Randelhoff et al. 2018), and high relative read abundance of phytoplankton, dominated by diatoms, *Phaeocystis pouchetii* and the genus *Micromonas* (Egge et al. 2021) indicating a spring bloom situation. The “bloom” dynamic of certain MALV ASVs suggests that while they are present in low abundance at other times of the year and at other depths, these ASVs are favored by the conditions in the surface during spring. This pattern may be due to an increased abundance of hosts, either blooming species, of e.g., diatoms or dinoflagellates, or grazers, such as copepods. The diatoms were dominated by *Chaetoceros socialis* and *Detonula confervacea* in May, which were replaced by a more diverse community in the August surface samples (Egge et al. 2021). A somewhat similar pattern was seen for dinoflagellates, with a few taxonomic groups dominating in May (unclassified Dinophyceae, *Gyrodinium* and *Woloszynskia*), and higher taxonomic richness in August. Furthermore, the dominating taxon in the metazoan metabarcoding reads shifted from the calanoid copepod genus *Calanus* in May, to the cyclopoid genus *Oithona* in August (Figure S4). Calanoid and cyclopoid copepod species have been shown to harbour different eukaryotic parasites and symbionts, and in particular, different MALV parasites (Savage et al. 2023). Another parasitic dinoflagellate genus, *Chytriodinium*, which is not a member of the MALV, but harbours known parasites of copepod eggs, was dominating the dinophyceae reads in the 50-200 *µ*m fraction in May, and had a strongly reduced abundance in August (Egge et al. 2021). Similar to our results, Clarke et al. (2019) found high abundance of a particular MALV operational taxonomic unit (OTU) associated with high Chl *a* values in the Antarctic ocean. However, this OTU was affiliated with DG-I-Clade 1, and thus did not correspond to any of the spring bloom-associated ASVs from the present study. Nevertheless, it is interesting that MALV genotypes from different subclades may dominate during periods of high phytoplankton biomass in different oceanographic regions.

**Figure 5.**
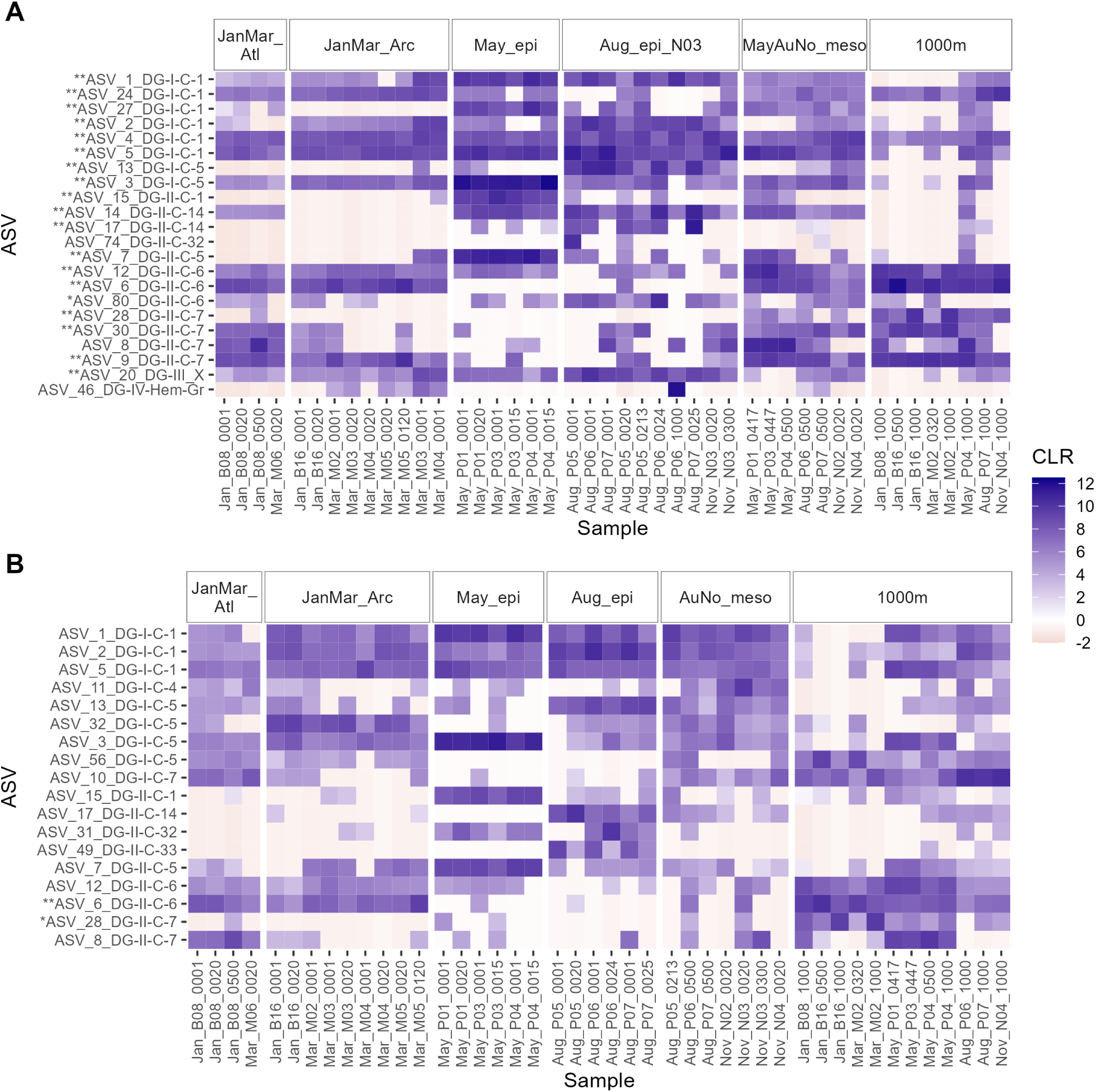
Distribution of the most abundant ASVs between sample clusters. The intensity of the colour indicates the centred log-ratio (CLR) of the ASV in a given sample. ASVs with CLR above 10 in at least one sample are included. (A) Pico (0.4-3 *µ*m) fraction, (B) nano-micro (3-200 *µ*m) fraction. * indicates whether an ASV has significantly different CLR between clusters. * p =< 0.1, ** p =< 0.05.

### Difference between the pico- and nano-micro fractions

MALV sequences from the smallest size fractions in environmental sequencing studies likely come from the dinospores (Guillou et al. 2008), but could also come from larger infected host cells that are disrupted during filtration. In addition to infecting larger hosts, dinospores may also be grazed by microzooplankton. Thus, in larger fractions obtained by sequential filtration, the presence of a MALV ASV may represent an infected host, dinospores recently ingested by microzooplankton, freeliving spores caught in the material on the filter, or potentially the vermiform stage. The spores are thought to be mainly short-lived, with a life span of 2-3 days (Coats and Park 2002), but it has been speculated that some MALV have evolved longer-living spores as an adaptation to life in a cold environment, as longer-living spores have been observed in other parasitic dinoflagellate groups (Coats 1999). If samples in the pico size fraction reflect recent spore releases from hosts in the nano-micro fraction, we would expect that the size fractions had the same taxonomic composition, possibly with different proportional abundance, and that the sample similarity relationships would be similar. This was mostly the case, as we observed similar clustering patterns in the pico- and nano-micro fractions (Figure 3). However, leakage between size fractions, as described above, cannot be ruled out. We may speculate that the ASVs and clades that are more abundant in the pico fraction, such as DG-II-Clade 6 and 7, represent spores released from hosts larger than 200 *µ*m.

### Distribution of phylogenetically similar ASVs

The composition of MALV ASVs in each sample cluster usually contained several clades from both Dino-Group I and II/IV (Figure S3). Previous studies have found that parasites of the same host type may come from several different clades within MALV (Guillou et al. 2008), e.g., both DG-I and DG-II/IV have members parasitizing other dinoflagellates. We also observed different distributions of ASVs within the same clade, e.g., ASV_1_DG-I-C1, which was relatively abundant in all samples, and ASV_25_DG-I-C1, which was only abundant in the Aug_epi cluster (Figure 6). There were also different distributions in the nano-micro fraction within DG-I-Clade 1 (ASV_171 - ASV_1, Figure 6 C). Some sister clades exhibited overall similar distributions, e.g., DG-II-Clades 6 and 7, which were abundant in the deep and dark samples. However, ASVs within these clades exhibited different distributions, e.g., ASV_6_DG-II-C6, which was abundant at 1000 m in the pico size fraction, vs. ASV_101_DG-II-C6, which was abundant in the JanMar_Arc cluster, but not detected at 1000 m. This illustrates a complex pattern of environment and host preference both within and between clades of MALV.

**Figure 6.**
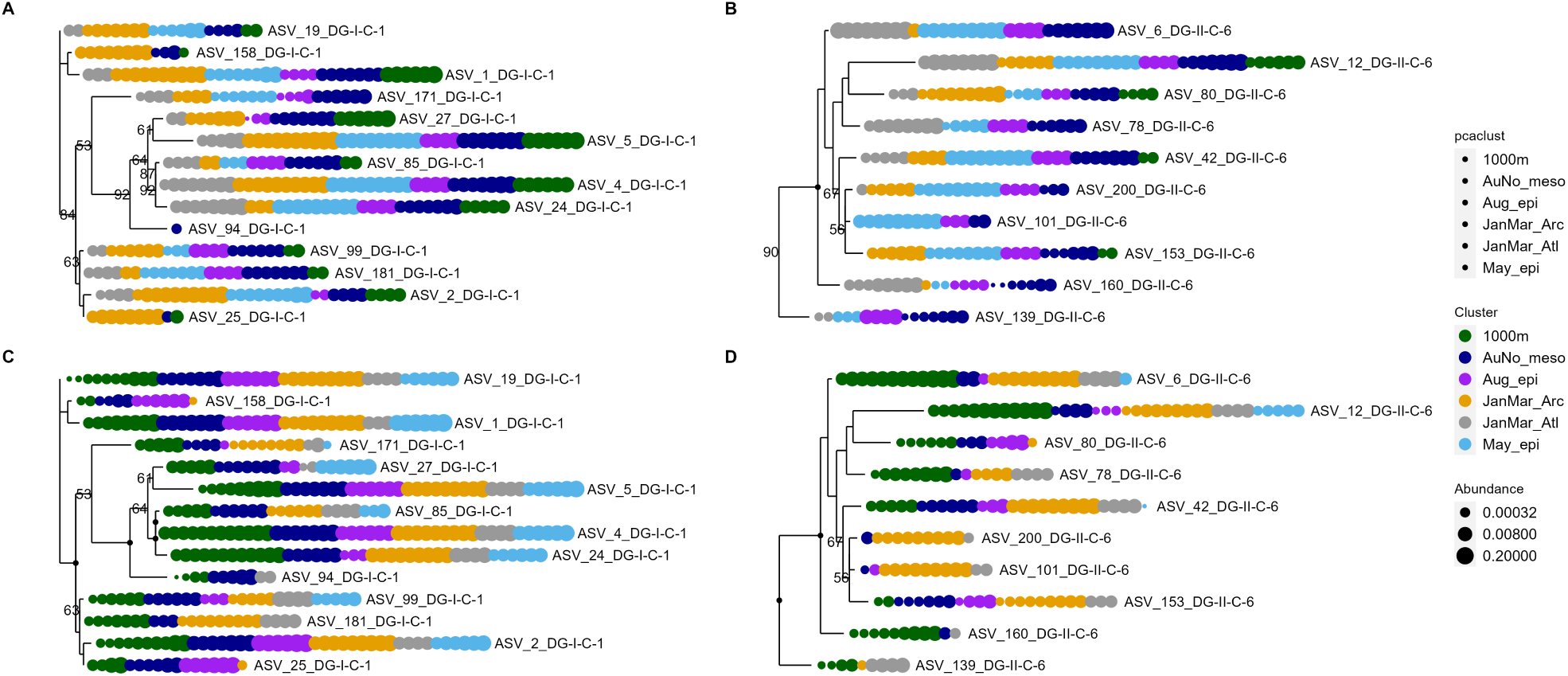
RAxML trees of the most abundant ASVs within the MALV clades Dino-Groups-I-Clade-1 and Dino-Group-II-Clade-6. (A) DG-I-Clade-1, pico fraction (B) DG-I-Clade-1, nano-micro fraction (C) DG-II-Clade-6, pico fraction (D) DG-II-Clade-6, nano-micro fraction. Each dot at the leaves corresponds to a sample. The size of the dot corresponds to relative abundance in the sample. Samples are colored according to cluster. Bootstrap values > 50 are shown on the nodes.

## Conclusions

Much work remains to be done to elucidate the impact of MALV parasitism in the marine microbial food web, in particular identifying host-parasite relationships and the extent of host specificity. Our study contributes to understanding the environmental preferences of several MALV clades and ASVs. Future work based on this study could include designing fluorescence *in situ* hybridisation probes based on the most abundant ASVs, and trying to detect them in potential hosts by epifluorescence microscopy. To further clarify the putative interactions between MALV and blooming photoautotrophs, we suggest targeting the ASVs dominating in the spring and summer surface samples. Identifying the hosts of these abundant MALV ASVs in the Arctic Ocean will contribute to quantifying the impact of parasitism on the Arctic food web.

## Acknowledgements

We would like to thank all members of the MicroPolar and CarbonBridge consortia, and the crews onboard R/V Lance and R/V Helmer Hanssen for their efforts during the sampling cruises.

## Author contributions statement

Conceptualization: EE, AL, DV, BE; Data curation: EE, DV; Funding aquisition: AL, BE; Investigation: EE, AL, BE; Methodology: EE, AL, BE; Project administration: AL, BE; Resources: AL; Supervision: DV, BE; Validation: EE, AL, DV, BE; Visualization: EE; Writing: EE, AL, DV, BE.

## Data Availablity

For availability of the original data, see Egge et al. (2021). ASV tables and R scripts from this study are deposited in GitHub at https://github.com/EEgge/micropolar_syndinialespaper.

## Supplementary Material

**Table S1.**
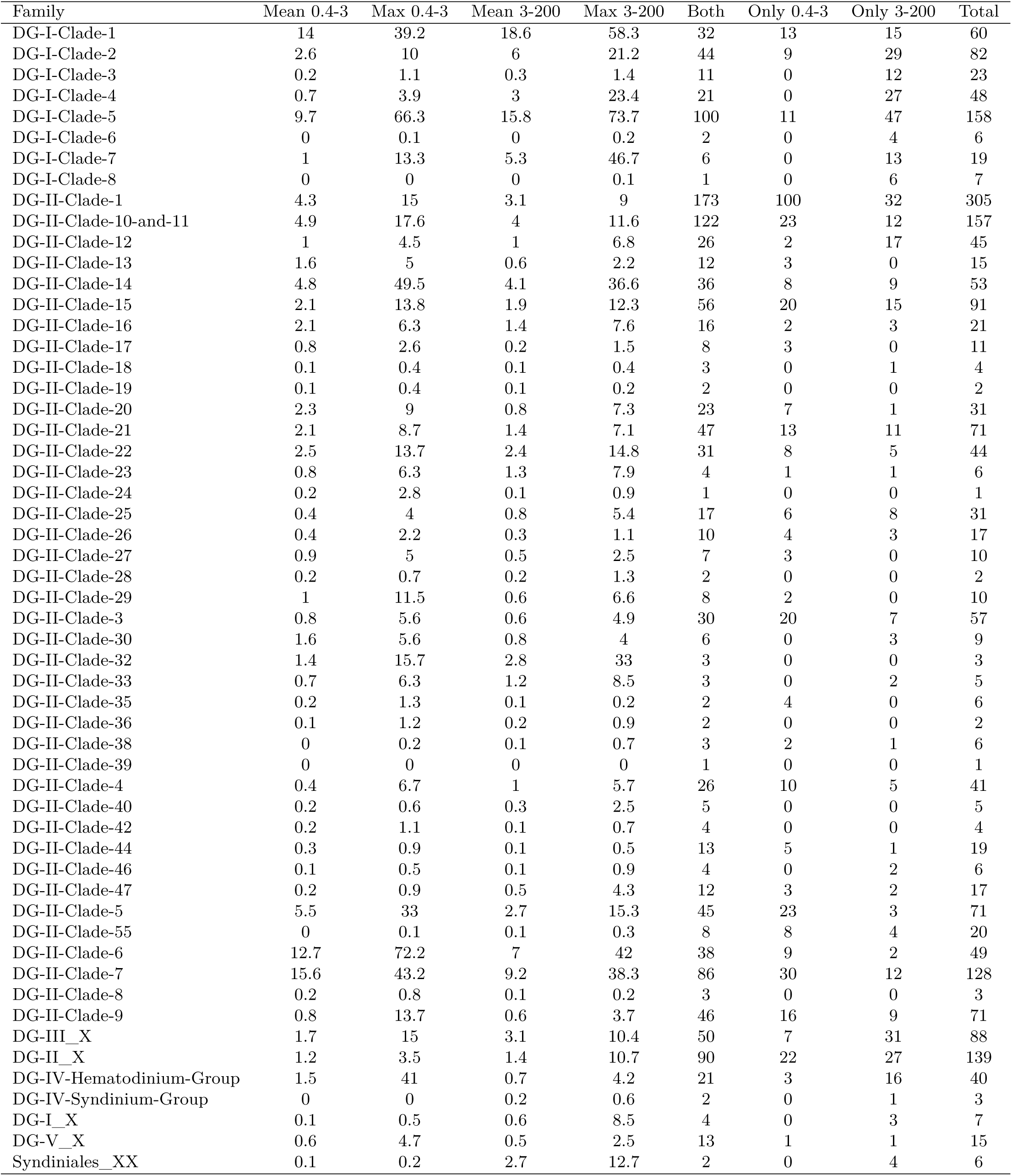
Table of MALV clades detected in the Micropolar 18S V4 metabarcoding data set. “Mean” = mean, “Max” = maximum percent read abundance in the pico- and nano-micro size fractions, respectively. “Both” = number of ASVs detected in both fractions, “Only” = number of ASVs only detected in each of the respective fractions, “Total” = the total number of ASVs assigned to each clade.

**Table S2.**
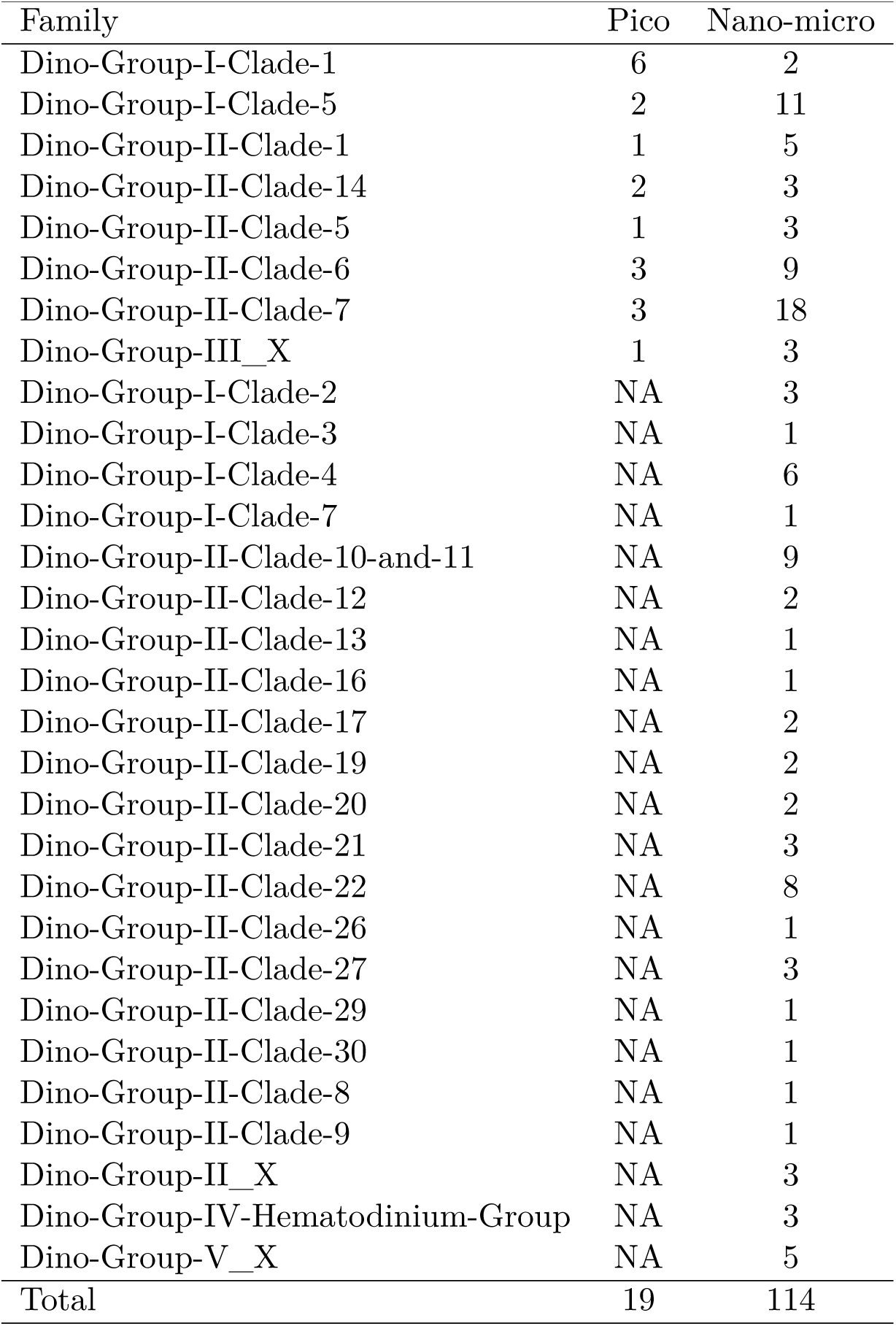
Number of ASVs per clade with significantly different centered log-ratio between clusters.

**Table S3.**
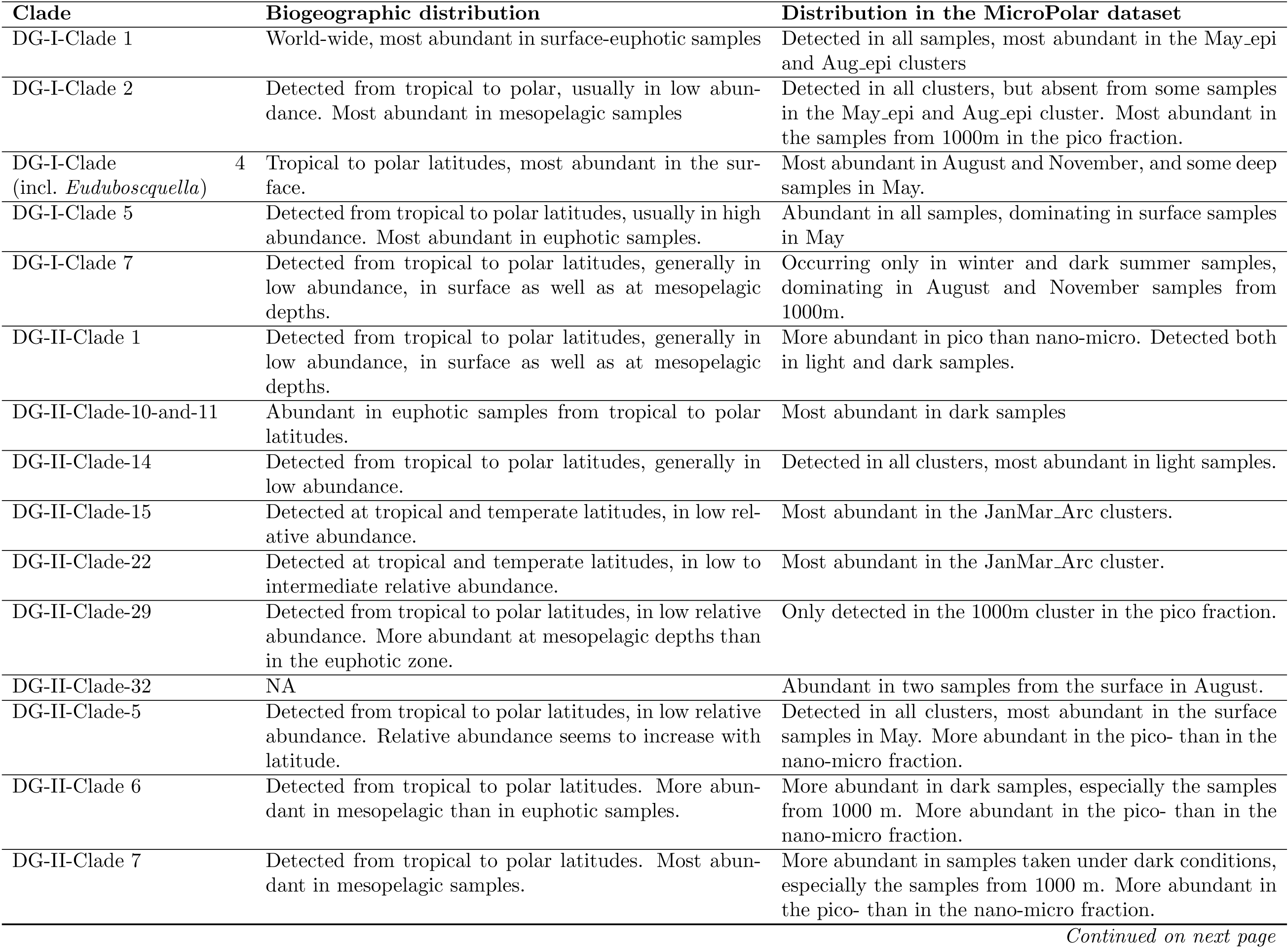
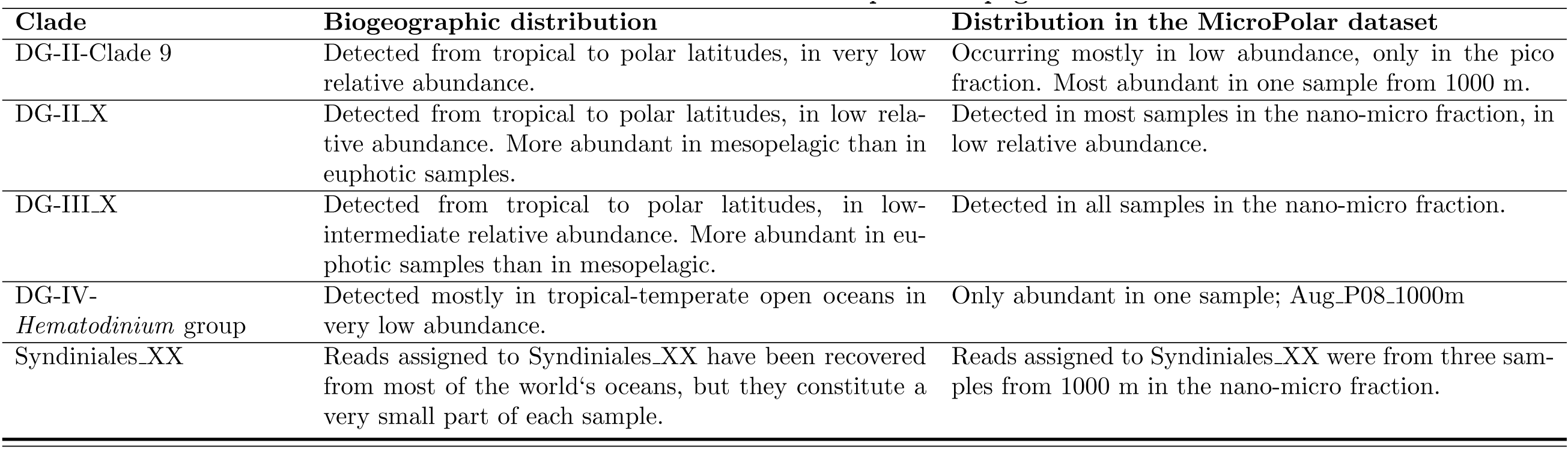
Biogeographic distribution of abundant MALV clades in the MicroPolar dataset, assessed by taxon search in metaPR^2.^

**Table S4.**
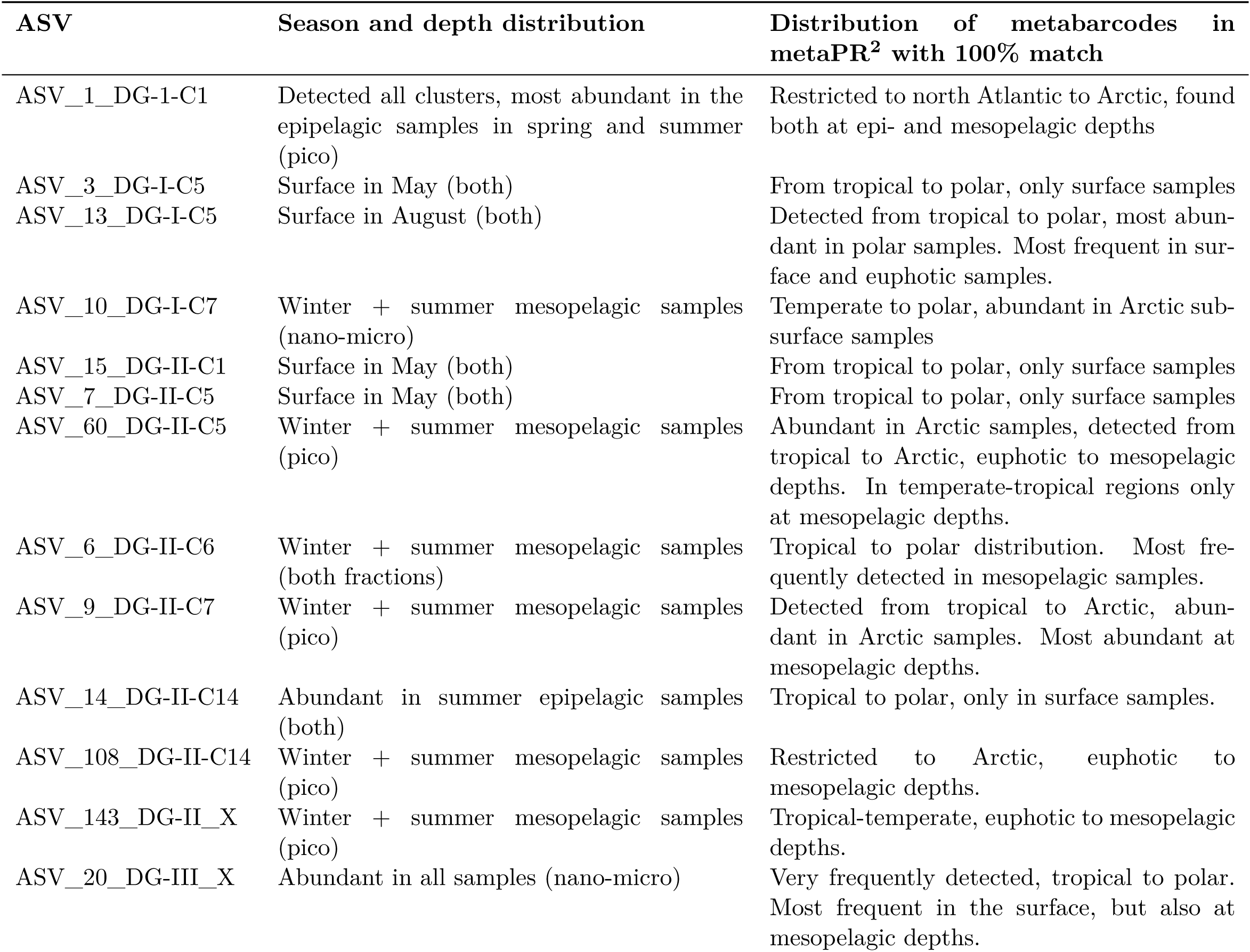
Biogeographic distribution of abundant MALV ASVs in the MicroPolar dataset, assessed by blast search in metaPR^2.^

**Figure S1.**
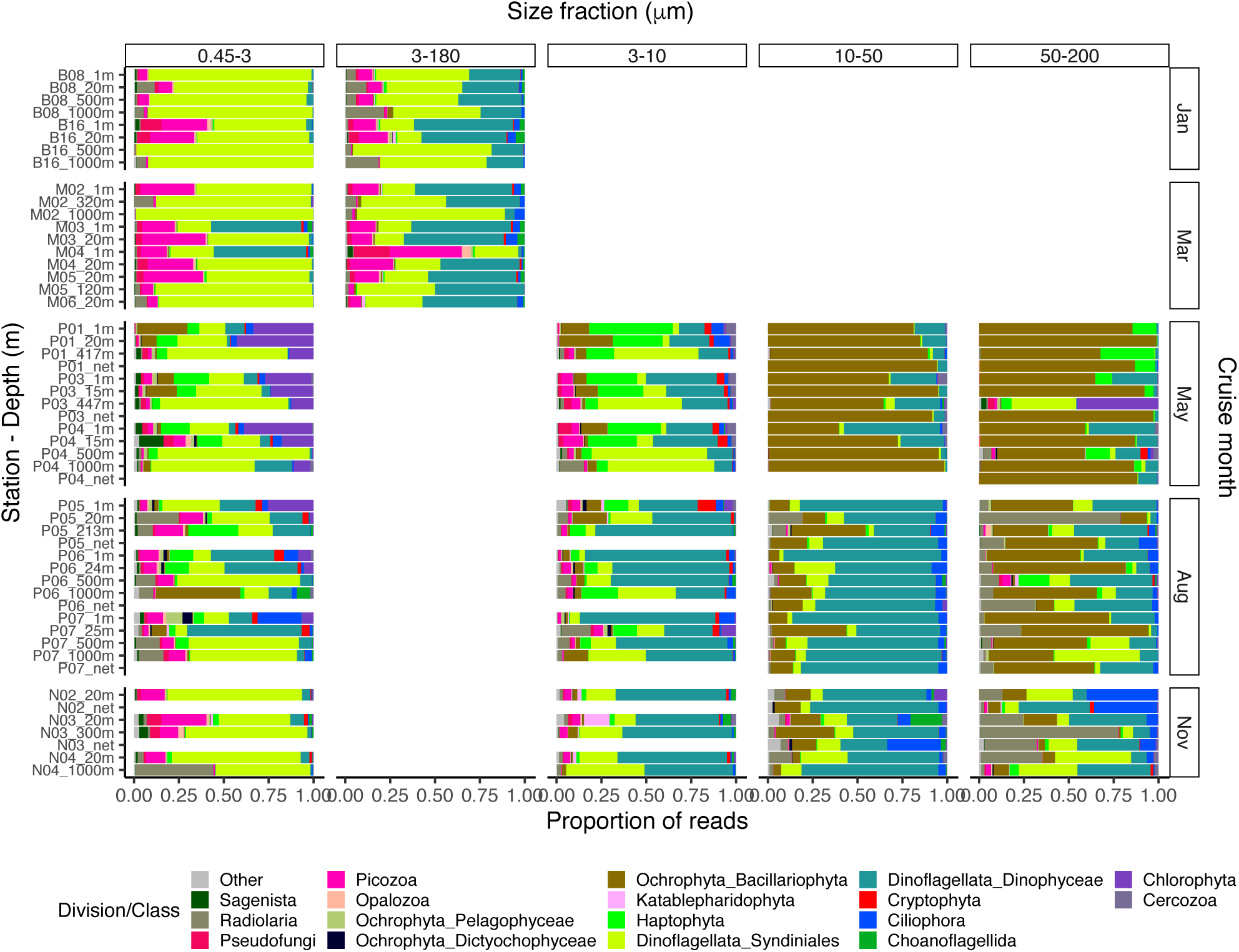
Relative abundance of the major protist groups in MicroPolar samples. Organised by sampling month (vertically) and size fraction (horizontally). MALV is highlighted in bright green.

**Figure S2.**
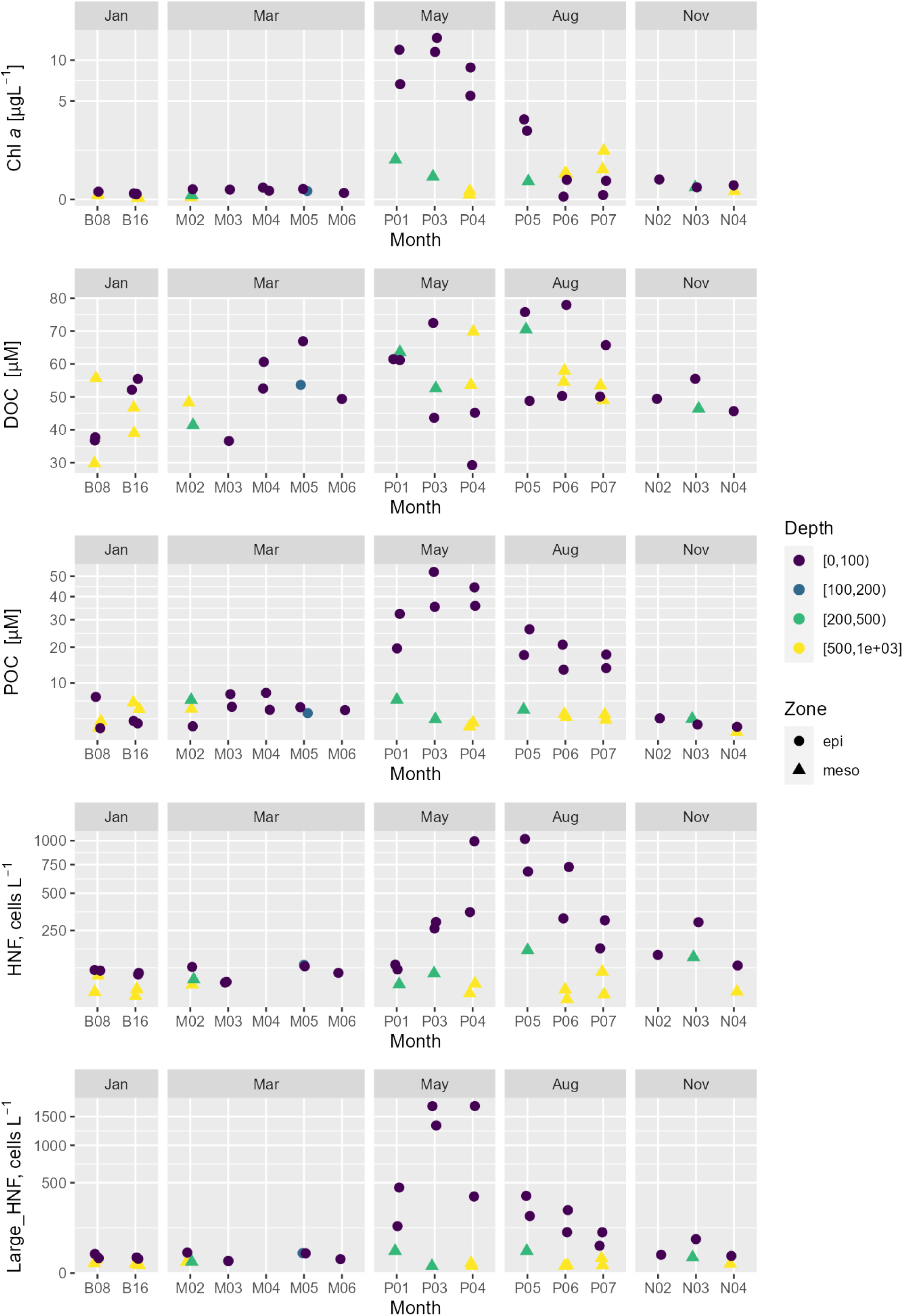
Components of organic matter in the water column. A) Chl *a*, B) and C) dissolved and particulate organic carbon (DOC, POC), D) and E) Small and large heterotrophic nanoflagellates (HNF)

**Figure S3.**
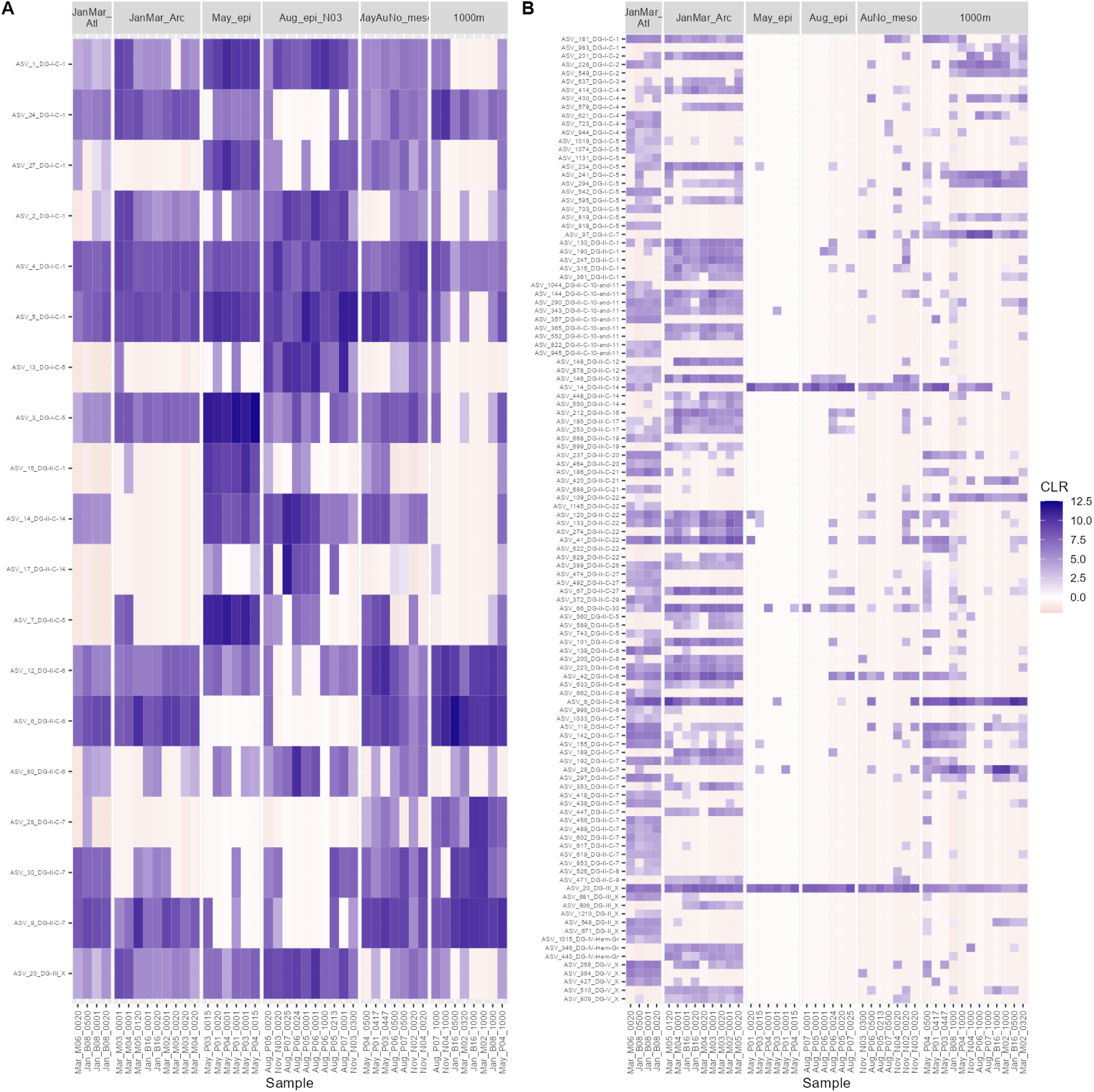
Heatmap of the centered log-ratio of ASVs with significant differential abundance between clusters. (A) pico-fraction (0.4-3 *µ*m), (B) nano-micro fraction (3-200 *µ*m)

**Figure S4.**
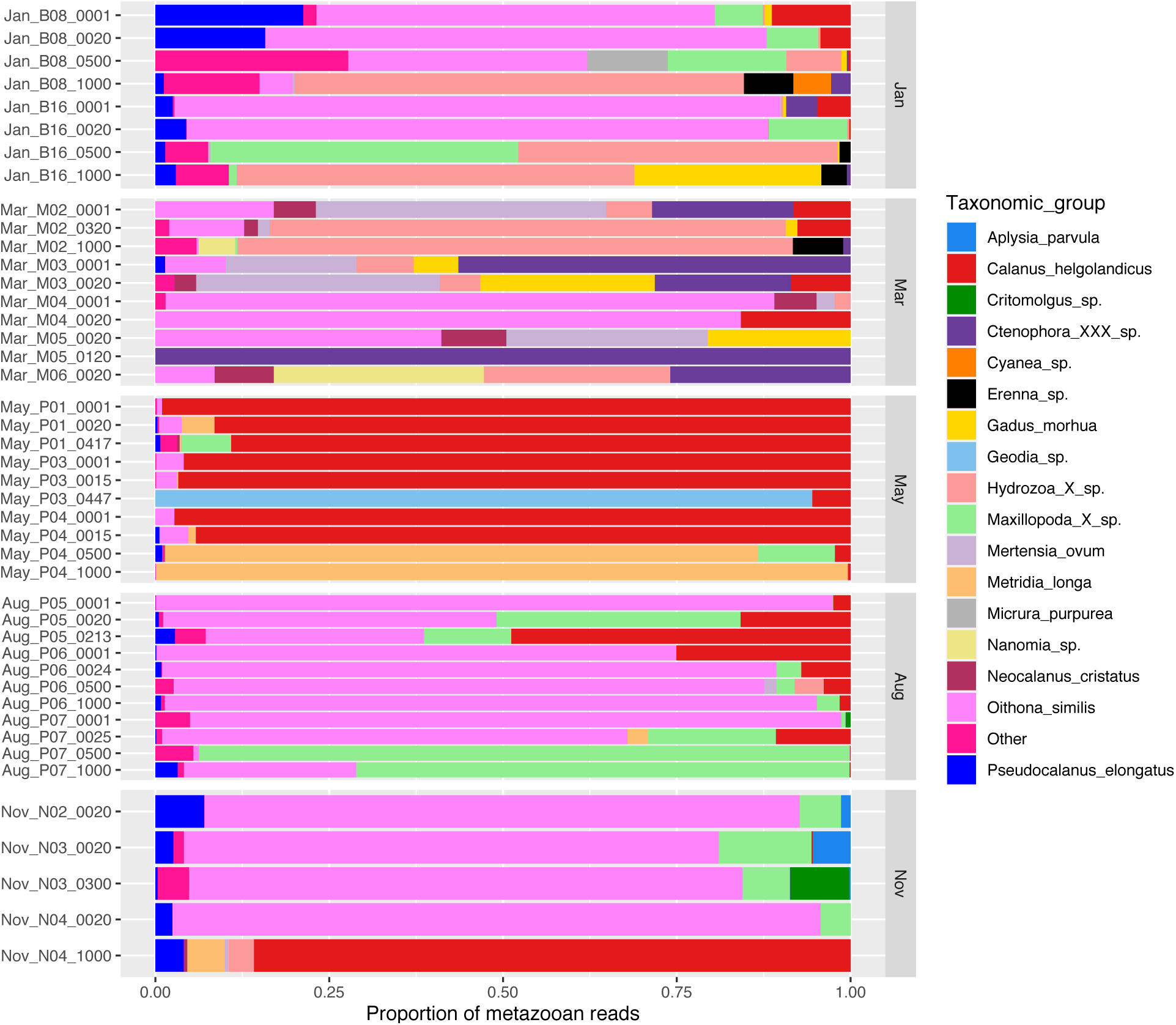
Taxonomic composition of metazoan reads. January and March: 3-180*µ*m fraction; May, August and November: 50-200*µ*m fraction.

**Figure S5.**
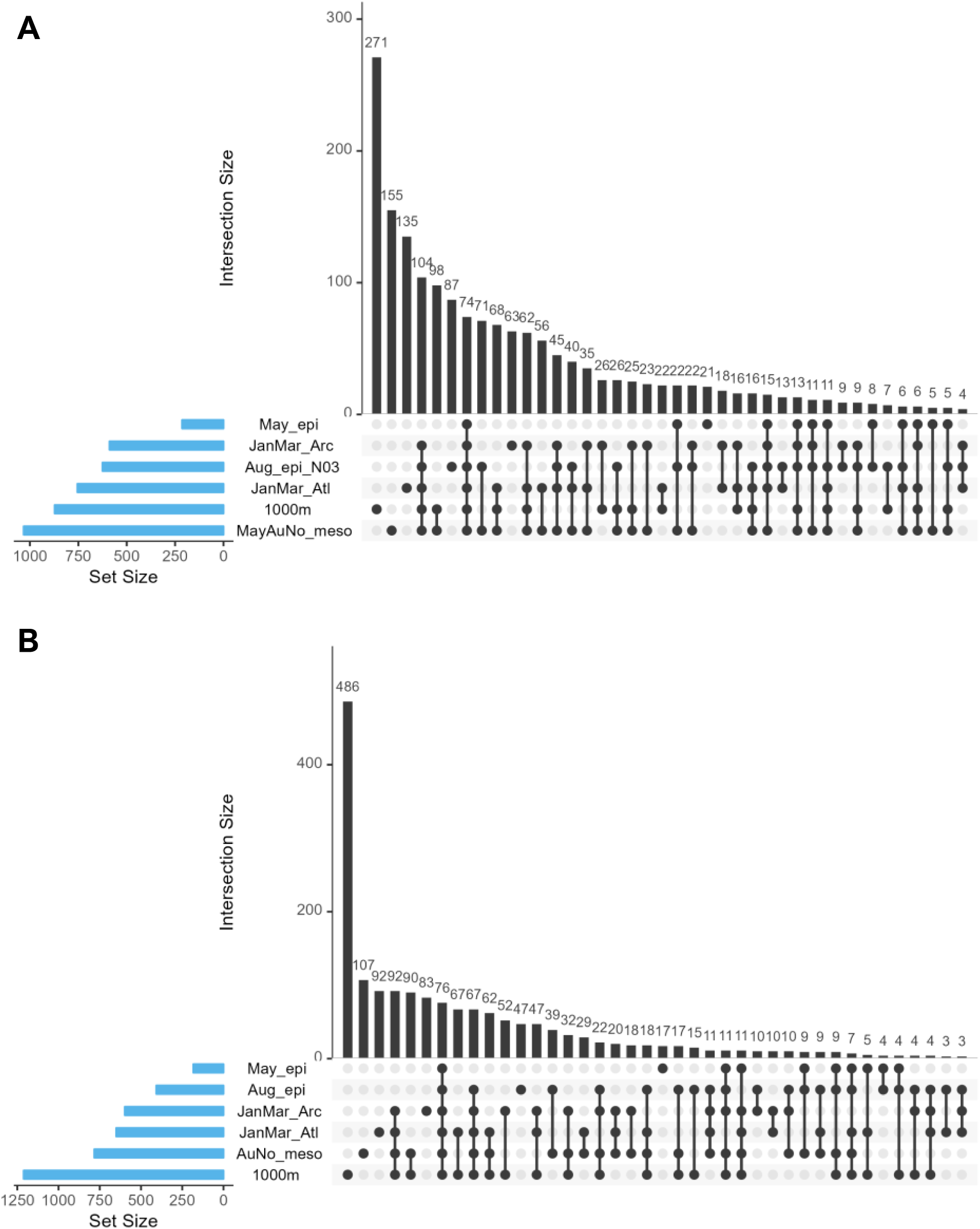
UpSet plots depicting the number of shared ASVs between sample clusters. (A) Shared ASVs in the pico (0.4-3 *µ*m) fraction, between the clusters of samples delimited in Figure 3. (B) Shared ASVs in the nano-micro (3-200 *µ*m) fraction.

## References cited

Anderson, S. R., L. Blanco-Bercial, C. A. Carlson, and E. L. Harvey (2024). The role of Syndiniales parasites in depth-specific networks and carbon flux in the oligotrophic ocean. *ISME Communications*, ycae014.

Anderson, S. R. and E. L. Harvey (2020). Temporal Variability and Ecological Interactions of Parasitic Marine Syndiniales in Coastal Protist Communities. mSphere 5.

Berdjeb, L., A. Parada, D. M. Needham, and J. A. Fuhrman (2018). Short-term dynamics and interactions of marine protist communities during the spring-summer transition. ISME Journal 12, 1907–1917.

Bjorbækmo, M. F. M., A. Evenstad, L. L. Røsæg, A. K. Krabberød, and R. Logares (2019). The planktonic protist interactome: where do we stand after a century of research? The ISME Journal, 1–16.

Borcard, D., F. Gillet, and P. Legendre (2018). Numerical Ecology with R. Use R!

Bråte, J., A. K. Krabberød, J. K. Dolven, R. F. Ose, T. Kristensen, K. R. Bjørklund, and K. Shalchian-Tabrizi (2012). Radiolaria Associated with Large Diversity of Marine Alveolates. Protist 163, 767– 777.

Cachon, J. (1964). Contribution a l’etude des Peridiniens parasites. Cytologie, cycles evolutifs. Ann. Sci. nat. zoologie, Paris 6, 1–158.

Callahan, B. J., P. J. McMurdie, M. J. Rosen, A. W. Han, A. J. A. Johnson, and S. P. Holmes (2016). DADA2: High-resolution sample inference from Illumina amplicon data. Nature Methods 13, 581– 583.

Capella-Gutiérrez, S., J. Silla-Martınez, and T. Gabaldón (n.d.). A tool for automated alignment trimming in large-scale phylogenetic analyses., 2009, 25. DOI: 10.1093/bioinformatics/btp348 (), 1972–1973.

Chambouvet, A., P. Morin, D. Marie, and L. Guillou (2008). Control of Toxic Marine Dinoflagellate Blooms by Serial Parasitic Killers. Science 322, 1254–1258.

Chatton, E. (1920). Les Peridiniens parasites : Morphologie, reproduction, ethologie. Arch. Zool. Exp. Gen. 59, 1–475.

Clarke, L. J., S. Bestley, A. Bissett, and B. E. Deagle (2019). A globally distributed Syndiniales parasite dominates the Southern Ocean micro-eukaryote community near the sea-ice edge. The ISME Journal 13, 734–737.

Cleary, A. C. and E. G. Durbin (2016). Unexpected prevalence of parasite 18S rDNA sequences in winter among Antarctic marine protists. *Journal of Plankton Research*, fbw005.

Coats, D. W. (1999). Parasitic life styles of marine dinoflagellates 1. Journal of Eukaryotic Microbiology 46, 402–409.

Coats, D. W. and M. G. Park (2002). Parasitism of photosynthetic dinoflagellates by three strains of *Amoebophrya* (Dinophyta): Parasite survival, infectivity, generation time, and host specificity. Journal of Phycology 38, 520–528.

Conway, J. R., A. Lex, and N. Gehlenborg (2017). UpSetR: An R package for the visualization of intersecting sets and their properties. Bioinformatics 33, 2938–2940.

Davies, C. E., J. E. Thomas, S. H. Malkin, F. M. Batista, A. F. Rowley, and C. J. Coates (2022). Hematodinium sp. infection does not drive collateral disease contraction in a crustacean host. Elife 11, e70356.

Drivdal, M., E. H. Kunisch, B. A. Bluhm, R. Gradinger, S. Falk-Petersen, and J. Berge (2021). Connections to the deep: Deep vertical migrations, an important part of the life cycle of Apherusa glacialis, an arctic ice-associated amphipod. Frontiers in Marine Science 8, 772766.

Egge, E., S. Elferink, D. Vaulot, U. John, G. Bratbak, A. Larsen, and B. Edvardsen (2021). An 18S V4 rRNA metabarcoding dataset of protist diversity in the Atlantic inflow to the Arctic Ocean, through the year and down to 1000 m depth. Earth System Science Data 13, 4913–4928.

Fu, Y., P. Zheng, X. Zhang, Q. Zhang, and D. Ji (2020). Protist Interactions and Seasonal Dynamics in the Coast of Yantai, Northern Yellow Sea of China as Revealed by Metabarcoding. Journal of Ocean University of China 2020 19:4 19, 961–974.

Gloor, G. B., J. M. Macklaim, V. Pawlowsky-Glahn, and J. J. Egozcue (2017). Microbiome datasets are compositional: And this is not optional. Frontiers in Microbiology 8, 1–6.

Groisillier, A., R. Massana, K. Valentin, D. Vaulot, and L. Guillou (2006). Genetic diversity and habitats of two enigmatic marine alveolate lineages. Aquatic Microbial Ecology 42, 277–291.

Guillou, L., M. Viprey, A. Chambouvet, R. M. Welsh, A. R. Kirkham, R. Massana, D. J. Scanlan, and A. Z. Worden (2008). Widespread occurrence and genetic diversity of marine parasitoids belonging to Syndiniales (Alveolata). Environmental Microbiology 10, 3349–3365.

Guillou, L. et al. (2013). The Protist Ribosomal Reference database (PR2): a catalog of unicellular eukaryote small sub-unit rRNA sequences with curated taxonomy. Nucleic acids research 41, D597– 604.

Hirche, H.-J. and B. Niehoff (1996). Reproduction of the Arctic copepod *Calanus hyperboreus* in the Greenland Sea-field and laboratory observations. Polar biology 16, 209–219.

Hobbs, L., N. S. Banas, F. R. Cottier, J. Berge, and M. Daase (2020). Eat or Sleep: Availability of Winter Prey Explains Mid-Winter and Spring Activity in an Arctic *Calanus* Population. Frontiers in Marine Science 7.

Holt, C. C., E. Hehenberger, D. V. Tikhonenkov, V. K. Jacko-Reynolds, N. Okamoto, E. C. Cooney, N. A. Irwin, and P. J. Keeling (2023). Multiple parallel origins of parasitic Marine Alveolates. Nature Communications 14, 7049.

Jackson, D. A. (1997). Compositional data in community ecology: the paradigm or peril of proportions? Ecology 78, 929–940.

Katoh, K., J. Rozewicki, and K. D. Yamada (2019). MAFFT online service: multiple sequence alignment, interactive sequence choice and visualization. Briefings in bioinformatics 20, 1160–1166.

Kimmerer, W. and A. McKinnon (1990). High mortality in a copepod population caused by a parasitic dinoflagellate. Marine Biology 107, 449–452.

Koid, A., W. C. Nelson, A. Mraz, and K. B. Heidelberg (2012). Comparative analysis of eukaryotic marine microbial assemblages from 18S rRNA gene or gene transcript clone libraries using different methods of extraction. Applied and environmental microbiology 78, 3958–3965.

Kunisch, E. H., B. A. Bluhm, M. Daase, R. Gradinger, H. Hop, I. A. Melnikov, Ø. Varpe, and J. Berge (2020). Pelagic occurrences of the ice amphipod Apherusa glacialis throughout the Arctic. Journal of Plankton Research 42, 73–86. eprint: https://academic.oup.com/plankt/article-pdf/42/1/73/32307201/fbz072.pdf.

Kvile, K. Ø., C. Ashjian, and R. Ji (2019). Pan-Arctic depth distribution of Diapausing *Calanus* copepods. The Biological Bulletin 237, 76–89.

Lahti, L. and S. Shetty (2012-2019). microbiome R package.

López-García, P., F. Rodríguez-Valera, C. Pedrós-Alió, and D. Moreira (2001). Unexpected diversity of small eukaryotes in deep-sea Antarctic plankton. Nature 409, 603–607.

Lovejoy, C., R. Massana, and C. Pedrós-Alió (2006). Diversity and distribution of marine microbial eukaryotes in the Arctic Ocean and adjacent seas. Applied and environmental microbiology 72, 3085–95.

Marquardt, M., A. Vader, E. I. Stübner, M. Reigstad, and T. M. Gabrielsen (2016). Strong seasonality of marine microbial eukaryotes in a high-Arctic fjord (Isfjorden, in West Spitsbergen, Norway). Applied and Environmental Microbiology 82, 1868–1880.

Massana, R., M. Pernice, J. a. Bunge, and J. del Campo (2011). Sequence diversity and novelty of natural assemblages of picoeukaryotes from the Indian Ocean. The ISME journal 5, 184–95.

McMurdie, P. J. and S. Holmes (2013). phyloseq: An R package for reproducible interactive analysis and graphics of microbiome census data. PLoS ONE 8, e61217.

McMurdie, P. J. and S. Holmes (2014). Waste not, want not: why rarefying microbiome data is inadmissible. PLoS computational biology 10, e1003531.

Nagata, T., C. Tamburini, J. Arıstegui, F. Baltar, A. B. Bochdansky, S. Fonda-Umani, H. Fukuda, A. Gogou, D. A. Hansell, R. L. Hansman, et al. (2010). Emerging concepts on microbial processes in the bathypelagic ocean–ecology, biogeochemistry, and genomics. Deep Sea Research Part II: Topical Studies in Oceanography 57, 1519–1536.

Nearing, J. T., G. M. Douglas, M. G. Hayes, J. MacDonald, D. K. Desai, N. Allward, C. M. Jones, R. J. Wright, A. S. Dhanani, A. M. Comeau, et al. (2022). Microbiome differential abundance methods produce different results across 38 datasets. Nature Communications 13, 342.

Oksanen, J., F. G. Blanchet, M. Friendly, R. Kindt, P. Legendre, D. McGlinn, P. Minchin, R. O’Hara, G. Simpson, P. Solymos, et al. (2021). vegan: Community Ecology Package. R package version 2.5-7. 2020.

Ollison, G. A., S. K. Hu, L. Y. Mesrop, E. F. DeLong, and D. A. Caron (2021). Come rain or shine: Depth not season shapes the active protistan community at station ALOHA in the North Pacific Subtropical Gyre. Deep Sea Research Part I: Oceanographic Research Papers 170, 103494.

Orr, R. J., S. A. Murray, A. Stüken, L. Rhodes, and K. S. Jakobsen (2012). When Naked Became Armored: An Eight-Gene Phylogeny Reveals Monophyletic Origin of Theca in Dinoflagellates. PLoS ONE 7, e50004.

Paulsen, M. L., H. Doré, L. Garczarek, L. Seuthe, O. Müller, R.-A. Sandaa, G. Bratbak, and A. Larsen (2016). Synechococcus in the Atlantic Gateway to the Arctic Ocean. Frontiers in Marine Science 3.

Pedersen, T. L. and M. Shemanarev (2022). ragg: Graphic Devices Based on AGG. R package version 1.2.2.

Pernice, M. C., C. R. Giner, R. Logares, J. Perera-Bel, S. G. Acinas, C. M. Duarte, J. M. Gasol, and R. Massana (2016). Large variability of bathypelagic microbial eukaryotic communities across the world’s oceans. The ISME Journal 10, 945–958.

Piredda, R., M. P. Tomasino, A. M. D’Erchia, C. Manzari, G. Pesole, M. Montresor, W. H. C. F. Kooistra, D. Sarno, and A. Zingone (2017). Diversity and temporal patterns of planktonic protist assemblages at a Mediterranean Long Term Ecological Research site. FEMS Microbiology Ecology 93, fiw200.

Poulin, R. and S. Morand (2000). The diversity of parasites. The quarterly review of biology 75, 277– 293.

Randelhoff, A., M. Reigstad, M. Chierici, A. Sundfjord, V. Ivanov, M. Cape, M. Vernet, J.-É. Tremblay, G. Bratbak, and S. Kristiansen (2018). Seasonality of the Physical and Biogeochemical Hydrography in the Inflow to the Arctic Ocean Through Fram Strait. Frontiers in Marine Science 5, 224.

Sandaa, R.-a., J. E. Storesund, E. Olesin, M. L. Paulsen, A. Larsen, G. Bratbak, and J. L. Ray (2018). Seasonality Drives Microbial Community Structure , Shaping both Eukaryotic and Prokaryotic Host – Viral Relationships in an Arctic Marine Ecosystem.

Sassenhagen, I., S. Irion, L. Jardillier, D. Moreira, and U. Christaki (2020). Protist interactions and community structure during early autumn in the Kerguelen region (Southern Ocean). Protist 171, 125709.

Savage, R.-L., J. L. Maud, C. T. E. Kellogg, B. P. V. Hunt, and V. Tai (2023). Symbiont diversity in the eukaryotic microbiomes of marine crustacean zooplankton. Journal of Plankton Research 45, 338–359. eprint: https://academic.oup.com/plankt/article-pdf/45/2/338/49720473/fbad003.pdf.

Sehein, T. R., R. J. Gast, M. Pachiadaki, L. Guillou, and V. P. Edgcomb (2022). Parasitic infections by Group II Syndiniales target selected dinoflagellate host populations within diverse protist assemblages in a model coastal pond. Environmental Microbiology 24, 1818–1834.

Seuthe, L., K. Rokkan Iversen, and F. Narcy (2011). Microbial processes in a high-latitude fjord (Kongsfjorden, Svalbard): II. Ciliates and dinoflagellates. Polar Biology 34, 751–766.

Skovgaard, A. (2014). Dirty tricks in the plankton: diversity and role of marine parasitic protists. Acta Protozoologica 53.

Skovgaard, A., R. Massana, V. Balague, and E. Saiz (2005). Phylogenetic position of the copepod-infesting parasite Syndinium turbo (Dinoflagellata, Syndinea). Protist 156, 413–423.

Strassert, J. F. H., A. Karnkowska, E. Hehenberger, J. del Campo, M. Kolisko, N. Okamoto, F. Burki, J. Janouškovec, C. Poirier, G. Leonard, S. J. Hallam, T. A. Richards, A. Z. Worden, A. E. Santoro, and P. J. Keeling (2018). Single cell genomics of uncultured marine alveolates shows paraphyly of basal dinoflagellates. The ISME Journal 12, 304–308.

Terrado, R., W. F. Vincent, and C. Lovejoy (2009). Mesopelagic protists: diversity and succession in a coastal Arctic ecosystem. Aquatic microbial ecology 56, 25–39.

Vargas, C. de, et al. (2015). Ocean plankton. Eukaryotic plankton diversity in the sunlit ocean. *Science (New York*, N.Y.) 348, 1261605.

Vaulot, D. (2022). pr2database. R package version 4.14.0.

Vaulot, D., C. W. H. Sim, D. Ong, B. Teo, C. Biwer, M. Jamy, and A. Lopes dos Santos (2022). metaPR2: a database of eukaryotic 18S rRNA metabarcodes with an emphasis on protists. Molecular Ecology Resources.

Wang, L.-G., T. T.-Y. Lam, S. Xu, Z. Dai, L. Zhou, T. Feng, P. Guo, C. W. Dunn, B. R. Jones, T. Bradley, H. Zhu, Y. Guan, Y. Jiang, and G. Yu (2020). treeio: an R package for phylogenetic tree input and output with richly annotated and associated data. Molecular Biology and Evolution 37, 599–603.

Wang, Q., G. M. Garrity, J. M. Tiedje, and J. R. Cole (2007). Naïve Bayesian Classifier for Rapid Assignment of rRNA Sequences into the New Bacterial Taxonomy. Applied and Environmental Microbiology 73, 5261–5267. eprint: https://journals.asm.org/doi/pdf/10.1128/AEM.00062-07.

Wang, Y., U. Naumann, D. Eddelbuettel, J. Wilshire, and D. Warton (2022). mvabund: Statistical Methods for Analysing Multivariate Abundance Data. R package version 4.2.1.

Wickham, H. (2016). ggplot2: Elegant Graphics for Data Analysis.

Wilson, B., O. Müller, E.-L. Nordmann, L. Seuthe, G. Bratbak, and L. Øvreås (2017). Changes in Marine Prokaryote Composition with Season and Depth Over an Arctic Polar Year. Frontiers in Marine Science 4, 1–17.

Worden, A. Z., M. J. Follows, S. J. Giovannoni, S. Wilken, A. E. Zimmerman, and P. J. Keeling (2015). Rethinking the marine carbon cycle: Factoring in the multifarious lifestyles of microbes. Science 347.

Xu, D., N. Jiao, R. Ren, and A. Warren (2017). Distribution and Diversity of Microbial Eukaryotes in Bathypelagic Waters of the South China Sea. Journal of Eukaryotic Microbiology 64, 370–382.

