## Supplemental material for "The community of Marine Alveolate parasites in the Atlantic inflow to the Arctic Ocean is structured by season, depth and water mass"

### Supplementary Material

**Table S1:** Table of MALV clades detected in the Micropolar 18S V4 metabarcoding data set. "Mean" = mean, "Max" = maximum percent read abundance in the pico- and nano-micro size fractions, respectively. "Both" = number of ASVs detected in both fractions, "Only" = number of ASVs only detected in each of the respective fractions, "Total" = the total number of ASVs assigned to each clade.

| Family | Mean 0.4-3 | Max 0.4-3 | Mean 3-200 | Max 3-200 | Both | Only 0.4-3 | Only 3-200 | Total |
|---|---|---|---|---|---|---|---|---|
| DG-I-Clade-1 | 14 | 39.2 | 18.6 | 58.3 | 32 | 13 | 15 | 60 |
| DG-I-Clade-2 | 2.6 | 10 | 6 | 21.2 | 44 | 9 | 29 | 82 |
| DG-I-Clade-3 | 0.2 | 1.1 | 0.3 | 1.4 | 11 | 0 | 12 | 23 |
| DG-I-Clade-4 | 0.7 | 3.9 | 3 | 23.4 | 21 | 0 | 27 | 48 |
| DG-I-Clade-5 | 9.7 | 66.3 | 15.8 | 73.7 | 100 | 11 | 47 | 158 |
| DG-I-Clade-6 | 0 | 0.1 | 0 | 0.2 | 2 | 0 | 4 | 6 |
| DG-I-Clade-7 | 1 | 13.3 | 5.3 | 46.7 | 6 | 0 | 13 | 19 |
| DG-I-Clade-8 | 0 | 0 | 0 | 0.1 | 1 | 0 | 6 | 7 |
| DG-II-Clade-1 | 4.3 | 15 | 3.1 | 9 | 173 | 100 | 32 | 305 |
| DG-II-Clade-10-and-11 | 4.9 | 17.6 | 4 | 11.6 | 122 | 23 | 12 | 157 |
| DG-II-Clade-12 | 1 | 4.5 | 1 | 6.8 | 26 | 2 | 17 | 45 |
| DG-II-Clade-13 | 1.6 | 5 | 0.6 | 2.2 | 12 | 3 | 0 | 15 |
| DG-II-Clade-14 | 4.8 | 49.5 | 4.1 | 36.6 | 36 | 8 | 9 | 53 |
| DG-II-Clade-15 | 2.1 | 13.8 | 1.9 | 12.3 | 56 | 20 | 15 | 91 |
| DG-II-Clade-16 | 2.1 | 6.3 | 1.4 | 7.6 | 16 | 2 | 3 | 21 |
| DG-II-Clade-17 | 0.8 | 2.6 | 0.2 | 1.5 | 8 | 3 | 0 | 11 |
| DG-II-Clade-18 | 0.1 | 0.4 | 0.1 | 0.4 | 3 | 0 | 1 | 4 |
| DG-II-Clade-19 | 0.1 | 0.4 | 0.1 | 0.2 | 2 | 0 | 0 | 2 |
| DG-II-Clade-20 | 2.3 | 9 | 0.8 | 7.3 | 23 | 7 | 1 | 31 |
| DG-II-Clade-21 | 2.1 | 8.7 | 1.4 | 7.1 | 47 | 13 | 11 | 71 |
| DG-II-Clade-22 | 2.5 | 13.7 | 2.4 | 14.8 | 31 | 8 | 5 | 44 |
| DG-II-Clade-23 | 0.8 | 6.3 | 1.3 | 7.9 | 4 | 1 | 1 | 6 |
| DG-II-Clade-24 | 0.2 | 2.8 | 0.1 | 0.9 | 1 | 0 | 0 | 1 |
| DG-II-Clade-25 | 0.4 | 4 | 0.8 | 5.4 | 17 | 6 | 8 | 31 |
| DG-II-Clade-26 | 0.4 | 2.2 | 0.3 | 1.1 | 10 | 4 | 3 | 17 |
| DG-II-Clade-27 | 0.9 | 5 | 0.5 | 2.5 | 7 | 3 | 0 | 10 |
| DG-II-Clade-28 | 0.2 | 0.7 | 0.2 | 1.3 | 2 | 0 | 0 | 2 |
| DG-II-Clade-29 | 1 | 11.5 | 0.6 | 6.6 | 8 | 2 | 0 | 10 |
| DG-II-Clade-3 | 0.8 | 5.6 | 0.6 | 4.9 | 30 | 20 | 7 | 57 |
| DG-II-Clade-30 | 1.6 | 5.6 | 0.8 | 4 | 6 | 0 | 3 | 9 |
| DG-II-Clade-32 | 1.4 | 15.7 | 2.8 | 33 | 3 | 0 | 0 | 3 |
| DG-II-Clade-33 | 0.7 | 6.3 | 1.2 | 8.5 | 3 | 0 | 2 | 5 |
| DG-II-Clade-35 | 0.2 | 1.3 | 0.1 | 0.2 | 2 | 4 | 0 | 6 |
| DG-II-Clade-36 | 0.1 | 1.2 | 0.2 | 0.9 | 2 | 0 | 0 | 2 |
| DG-II-Clade-38 | 0 | 0.2 | 0.1 | 0.7 | 3 | 2 | 1 | 6 |
| DG-II-Clade-39 | 0 | 0 | 0 | 0 | 1 | 0 | 0 | 1 |
| DG-II-Clade-4 | 0.4 | 6.7 | 1 | 5.7 | 26 | 10 | 5 | 41 |
| DG-II-Clade-40 | 0.2 | 0.6 | 0.3 | 2.5 | 5 | 0 | 0 | 5 |
| DG-II-Clade-42 | 0.2 | 1.1 | 0.1 | 0.7 | 4 | 0 | 0 | 4 |
| DG-II-Clade-44 | 0.3 | 0.9 | 0.1 | 0.5 | 13 | 5 | 1 | 19 |
| DG-II-Clade-46 | 0.1 | 0.5 | 0.1 | 0.9 | 4 | 0 | 2 | 6 |
| DG-II-Clade-47 | 0.2 | 0.9 | 0.5 | 4.3 | 12 | 3 | 2 | 17 |
| DG-II-Clade-5 | 5.5 | 33 | 2.7 | 15.3 | 45 | 23 | 3 | 71 |
| DG-II-Clade-55 | 0 | 0.1 | 0.1 | 0.3 | 8 | 8 | 4 | 20 |
| DG-II-Clade-6 | 12.7 | 72.2 | 7 | 42 | 38 | 9 | 2 | 49 |
| DG-II-Clade-7 | 15.6 | 43.2 | 9.2 | 38.3 | 86 | 30 | 12 | 128 |
| DG-II-Clade-8 | 0.2 | 0.8 | 0.1 | 0.2 | 3 | 0 | 0 | 3 |
| DG-II-Clade-9 | 0.8 | 13.7 | 0.6 | 3.7 | 46 | 16 | 9 | 71 |
| DG-III__X | 1.7 | 15 | 3.1 | 10.4 | 50 | 7 | 31 | 88 |
| DG-II__X | 1.2 | 3.5 | 1.4 | 10.7 | 90 | 22 | 27 | 139 |
| DG-IV-Hematodinium-Group | 1.5 | 41 | 0.7 | 4.2 | 21 | 3 | 16 | 40 |
| DG-IV-Syndinium-Group | 0 | 0 | 0.2 | 0.6 | 2 | 0 | 1 | 3 |
| DG-I__X | 0.1 | 0.5 | 0.6 | 8.5 | 4 | 0 | 3 | 7 |
| DG-V__X | 0.6 | 4.7 | 0.5 | 2.5 | 13 | 1 | 1 | 15 |
| Syndiniales__XX | 0.1 | 0.2 | 2.7 | 12.7 | 2 | 0 | 4 | 6 |

**Table S2:** Number of ASVs per clade with significantly different centered log-ratio between clusters.

| Family | Pico | Nano-micro |
|---|---|---|
| Dino-Group-I-Clade-1 | 6 | 2 |
| Dino-Group-I-Clade-5 | 2 | 11 |
| Dino-Group-II-Clade-1 | 1 | 5 |
| Dino-Group-II-Clade-14 | 2 | 3 |
| Dino-Group-II-Clade-5 | 1 | 3 |
| Dino-Group-II-Clade-6 | 3 | 9 |
| Dino-Group-II-Clade-7 | 3 | 18 |
| Dino-Group-III_X | 1 | 3 |
| Dino-Group-I-Clade-2 | NA | 3 |
| Dino-Group-I-Clade-3 | NA | 1 |
| Dino-Group-I-Clade-4 | NA | 6 |
| Dino-Group-I-Clade-7 | NA | 1 |
| Dino-Group-II-Clade-10-and-11 | NA | 9 |
| Dino-Group-II-Clade-12 | NA | 2 |
| Dino-Group-II-Clade-13 | NA | 1 |
| Dino-Group-II-Clade-16 | NA | 1 |
| Dino-Group-II-Clade-17 | NA | 2 |
| Dino-Group-II-Clade-19 | NA | 2 |
| Dino-Group-II-Clade-20 | NA | 2 |
| Dino-Group-II-Clade-21 | NA | 3 |
| Dino-Group-II-Clade-22 | NA | 8 |
| Dino-Group-II-Clade-26 | NA | 1 |
| Dino-Group-II-Clade-27 | NA | 3 |
| Dino-Group-II-Clade-29 | NA | 1 |
| Dino-Group-II-Clade-30 | NA | 1 |
| Dino-Group-II-Clade-8 | NA | 1 |
| Dino-Group-II-Clade-9 | NA | 1 |
| Dino-Group-II_X | NA | 3 |
| Dino-Group-IV-Hematodinium-Group | NA | 3 |
| Dino-Group-V_X | NA | 5 |
| Total | 19 | 114 |

**Table S3:** Biogeographic distribution of abundant MALV clades in the MicroPolar dataset, assessed by taxon search in metaPR$^2$.

| Clade | Biogeographic distribution | Distribution in the MicroPolar dataset |
|---|---|---|
| DG-I-Clade 1 | World-wide, most abundant in surface-euphotic samples | Detected in all samples, most abundant in the May_epi and Aug_epi clusters |
| DG-I-Clade 2 | Detected from tropical to polar, usually in low abundance. Most abundant in mesopelagic samples | Detected in all clusters, but absent from some samples in the May_epi and Aug_epi cluster. Most abundant in the samples from 1000m in the pico fraction. |
| DG-I-Clade 4 (incl. *Euduboscquella*) | Tropical to polar latitudes, most abundant in the surface. | Most abundant in August and November, and some deep samples in May. |
| DG-I-Clade 5 | Detected from tropical to polar latitudes, usually in high abundance. Most abundant in euphotic samples. | Abundant in all samples, dominating in surface samples in May |
| DG-I-Clade 7 | Detected from tropical to polar latitudes, generally in low abundance, in surface as well as at mesopelagic depths. | Occurring only in winter and dark summer samples, dominating in August and November samples from 1000m. |
| DG-II-Clade 1 | Detected from tropical to polar latitudes, generally in low abundance, in surface as well as at mesopelagic depths. | More abundant in pico than nano-micro. Detected both in light and dark samples. |
| DG-II-Clade-10-and-11 | Abundant in euphotic samples from tropical to polar latitudes. | Most abundant in dark samples |
| DG-II-Clade-14 | Detected from tropical to polar latitudes, generally in low abundance. | Detected in all clusters, most abundant in light samples. |
| DG-II-Clade-15 | Detected at tropical and temperate latitudes, in low relative abundance. | Most abundant in the JanMar_Arc clusters. |
| DG-II-Clade-22 | Detected at tropical and temperate latitudes, in low to intermediate relative abundance. | Most abundant in the JanMar_Arc cluster. |
| DG-II-Clade-29 | Detected from tropical to polar latitudes, in low relative abundance. More abundant at mesopelagic depths than in the euphotic zone. | Only detected in the 1000m cluster in the pico fraction. |
| DG-II-Clade-32 | NA | Abundant in two samples from the surface in August. |
| DG-II-Clade-5 | Detected from tropical to polar latitudes, in low relative abundance. Relative abundance seems to increase with latitude. | Detected in all clusters, most abundant in the surface samples in May. More abundant in the pico- than in the nano-micro fraction. |
| DG-II-Clade 6 | Detected from tropical to polar latitudes. More abundant in mesopelagic than in euphotic samples. | More abundant in dark samples, especially the samples from 1000 m. More abundant in the pico- than in the nano-micro fraction. |
| DG-II-Clade 7 | Detected from tropical to polar latitudes. Most abundant in mesopelagic samples. | More abundant in samples taken under dark conditions, especially the samples from 1000 m. More abundant in the pico- than in the nano-micro fraction. |

*Continued on next page*

**Table S3 – continued from previous page**

| Clade | Biogeographic distribution | Distribution in the MicroPolar dataset |
|---|---|---|
| DG-II-Clade 9 | Detected from tropical to polar latitudes, in very low relative abundance. | Occurring mostly in low abundance, only in the pico fraction. Most abundant in one sample from 1000 m. |
| DG-II_X | Detected from tropical to polar latitudes, in low relative abundance. More abundant in mesopelagic than in euphotic samples. | Detected in most samples in the nano-micro fraction, in low relative abundance. |
| DG-III_X | Detected from tropical to polar latitudes, in low-intermediate relative abundance. More abundant in euphotic samples than in mesopelagic. | Detected in all samples in the nano-micro fraction. |
| DG-IV-*Hematodinium* group | Detected mostly in tropical-temperate open oceans in very low abundance. | Only abundant in one sample; Aug_P08_1000m |
| Syndiniales_XX | Reads assigned to Syndiniales_XX have been recovered from most of the world‘s oceans, but they constitute a very small part of each sample. | Reads assigned to Syndiniales_XX were from three samples from 1000 m in the nano-micro fraction. |

**Table S4:** Biogeographic distribution of abundant MALV ASVs in the MicroPolar dataset, assessed by blast search in metaPR$^2$.

| ASV | Season and depth distribution | Distribution of metabarcodes in metaPR$^2$ with 100% match |
|---|---|---|
| ASV_1_DG-1-C1 | Detected all clusters, most abundant in the epipelagic samples in spring and summer (pico) | Restricted to north Atlantic to Arctic, found both at epi- and mesopelagic depths |
| ASV_3_DG-I-C5 | Surface in May (both) | From tropical to polar, only surface samples |
| ASV_13_DG-I-C5 | Surface in August (both) | Detected from tropical to polar, most abundant in polar samples. Most frequent in surface and euphotic samples. |
| ASV_10_DG-I-C7 | Winter + summer mesopelagic samples (nano-micro) | Temperate to polar, abundant in Arctic subsurface samples |
| ASV_15_DG-II-C1 | Surface in May (both) | From tropical to polar, only surface samples |
| ASV_7_DG-II-C5 | Surface in May (both) | From tropical to polar, only surface samples |
| ASV_60_DG-II-C5 | Winter + summer mesopelagic samples (pico) | Abundant in Arctic samples, detected from tropical to Arctic, euphotic to mesopelagic depths. In temperate-tropical regions only at mesopelagic depths. |
| ASV_6_DG-II-C6 | Winter + summer mesopelagic samples (both fractions) | Tropical to polar distribution. Most frequently detected in mesopelagic samples. |
| ASV_9_DG-II-C7 | Winter + summer mesopelagic samples (pico) | Detected from tropical to Arctic, abundant in Arctic samples. Most abundant at mesopelagic depths. |
| ASV_14_DG-II-C14 | Abundant in summer epipelagic samples (both) | Tropical to polar, only in surface samples. |
| ASV_108_DG-II-C14 | Winter + summer mesopelagic samples (pico) | Restricted to Arctic, euphotic to mesopelagic depths. |
| ASV_143_DG-II_X | Winter + summer mesopelagic samples (pico) | Tropical-temperate, euphotic to mesopelagic depths. |
| ASV_20_DG-III_X | Abundant in all samples (nano-micro) | Very frequently detected, tropical to polar. Most frequent in the surface, but also at mesopelagic depths. |

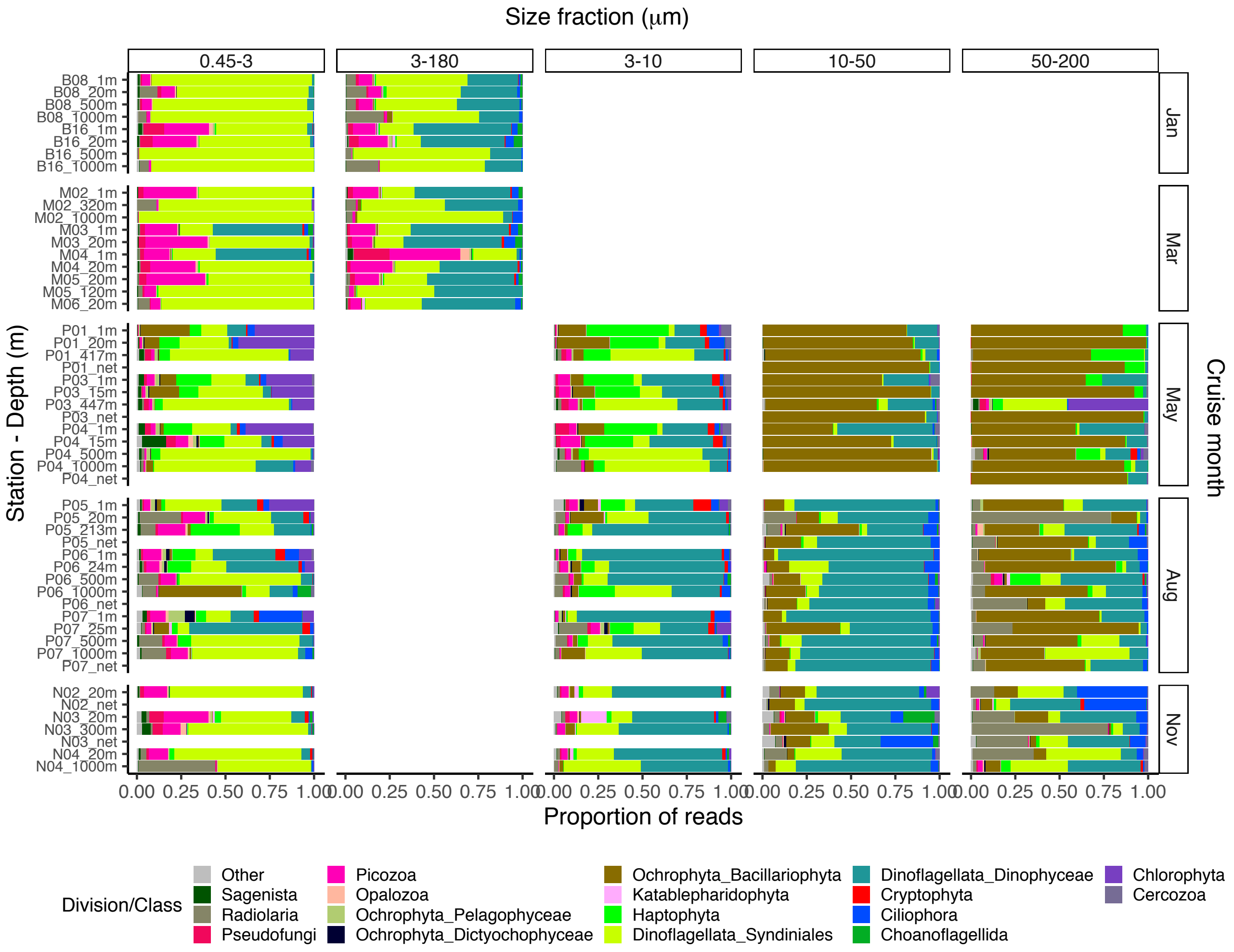


**Figure S1: Relative abundance of the major protist groups in MicroPolar samples.** Organised by sampling month (vertically) and size fraction (horizontally). MALV is highlighted in bright green.

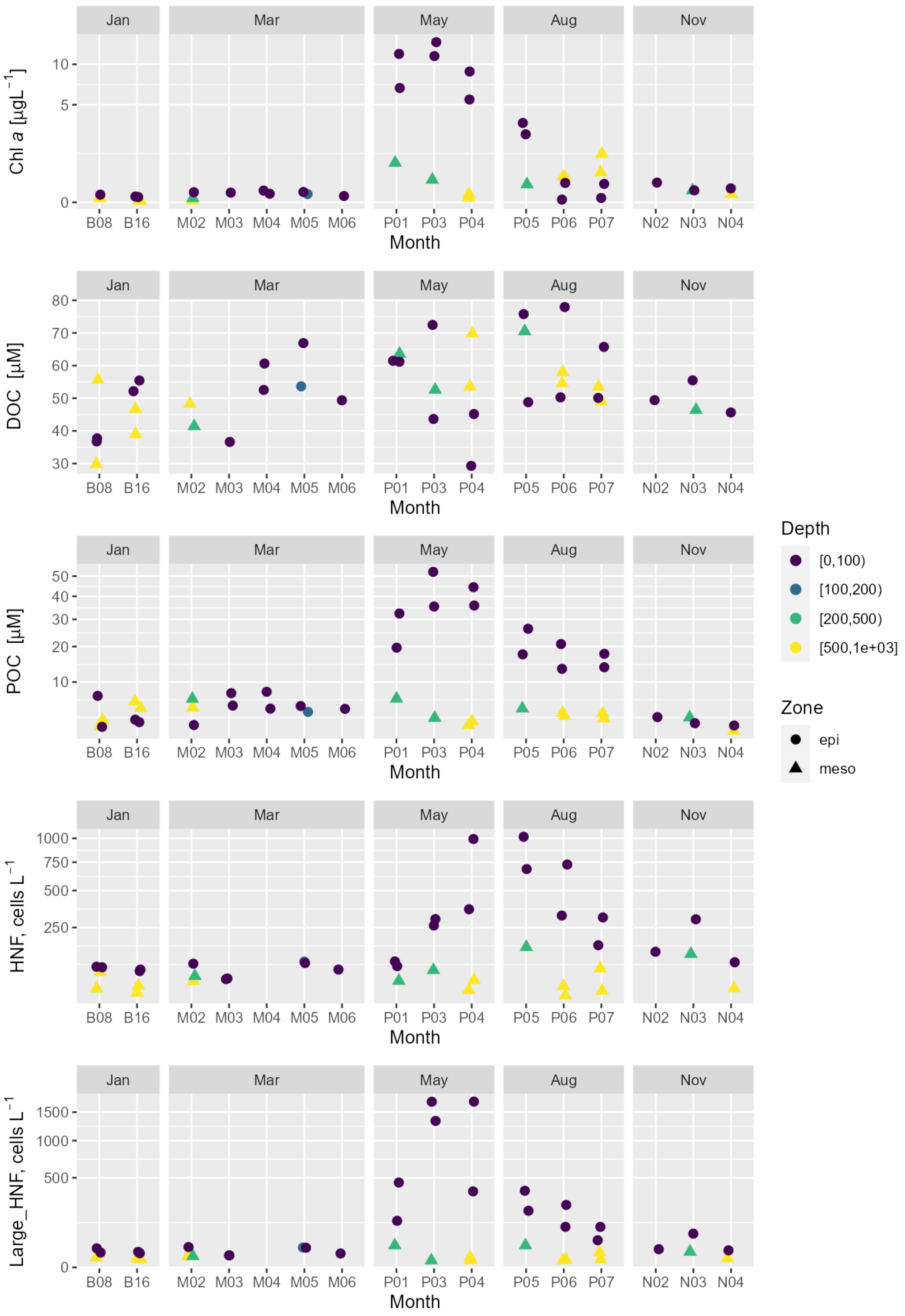


**Figure S2: Components of organic matter in the water column.** A) Chl *a*, B) and C) dissolved and particulate organic carbon (DOC, POC), D) and E) Small and large heterotrophic nanoflagellates (HNF)

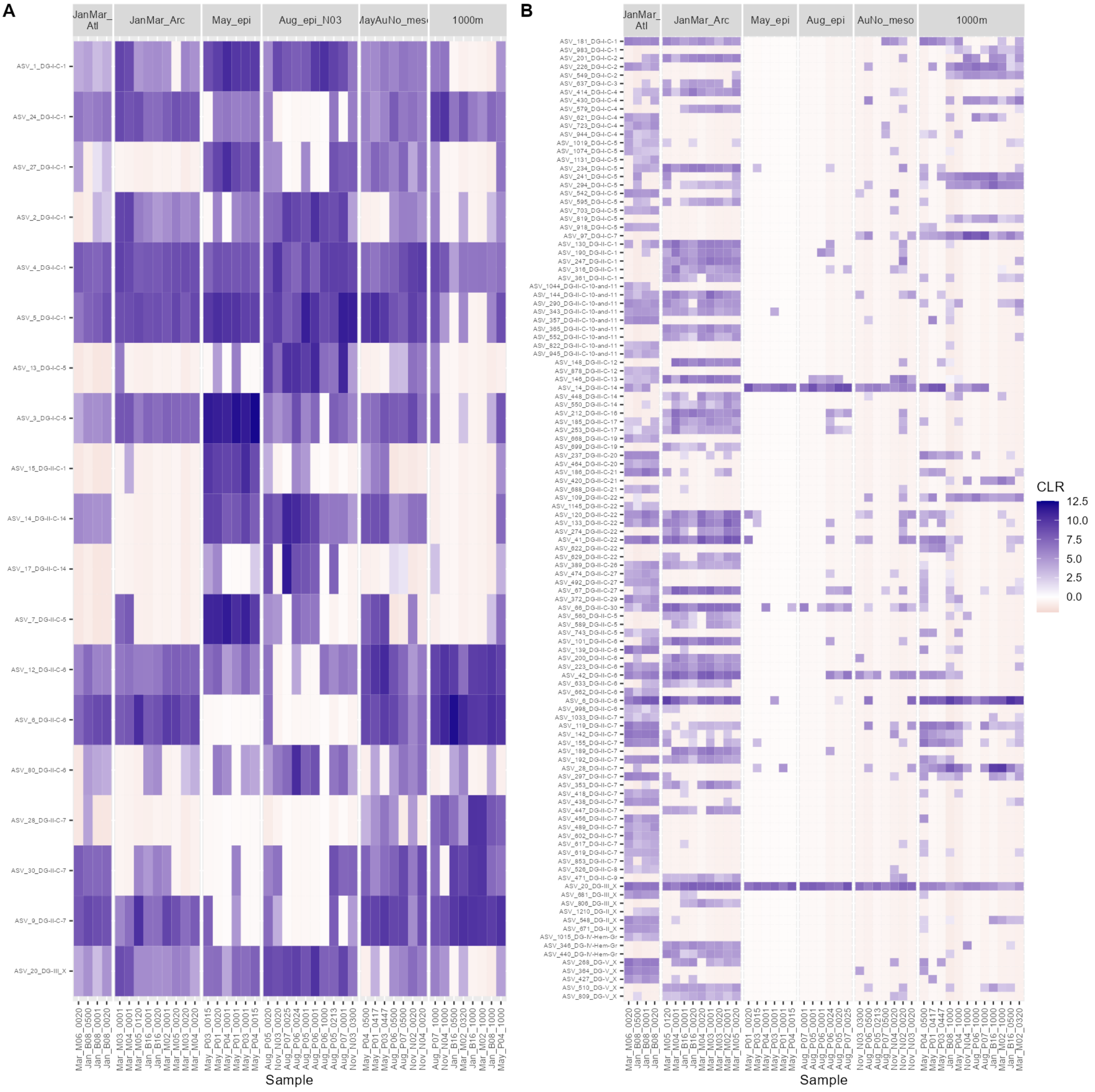


**Figure S3: Heatmap of the centered log-ratio of ASVs with significant differential abundance between clusters.** (A) pico-fraction (0.4-3 $\mu$m), (B) nano-micro fraction (3-200 $\mu$m)

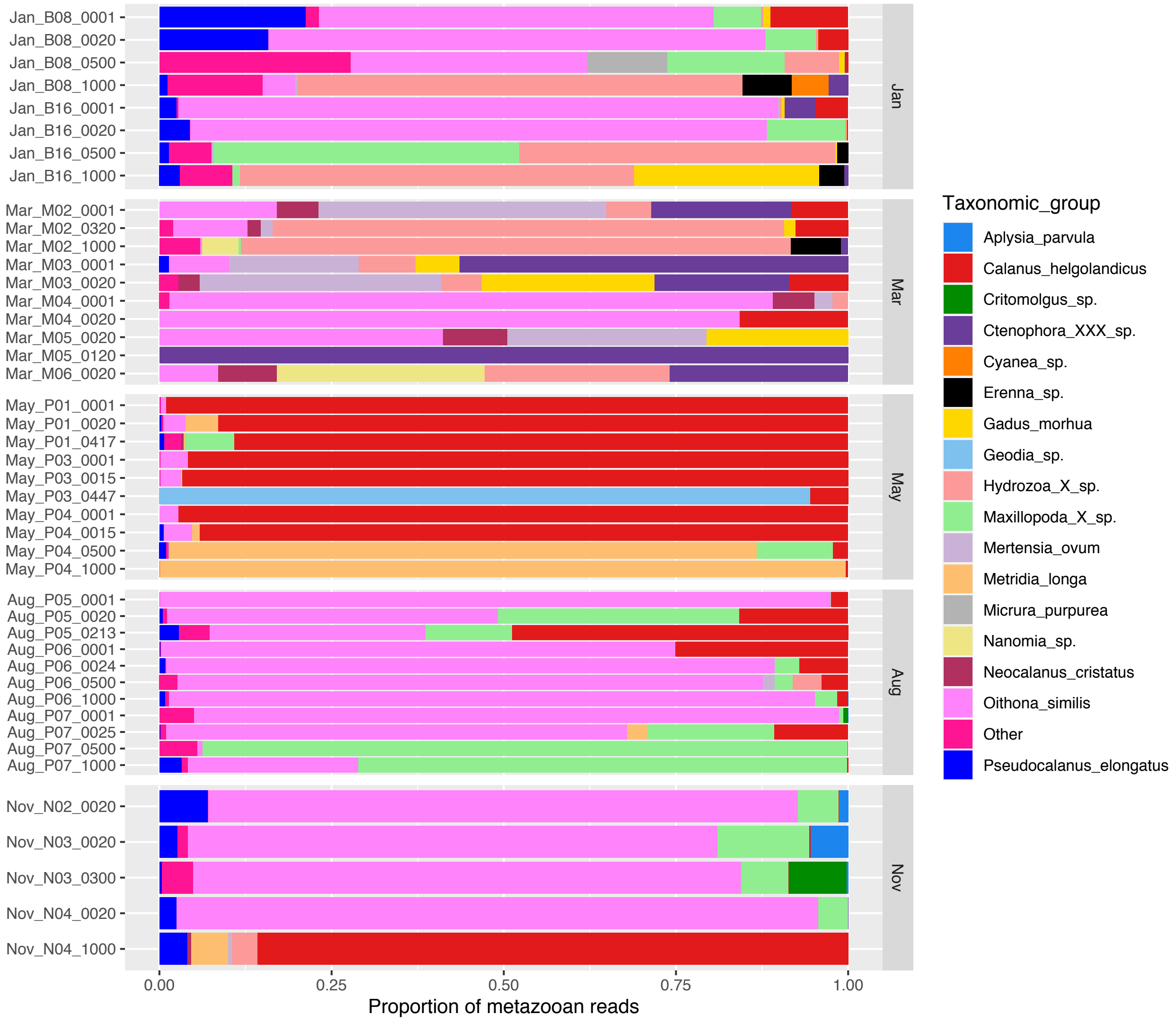


**Figure S4: Taxonomic composition of metazoan reads.** January and March: 3-180$\mu$m fraction; May, August and November: 50-200$\mu$m fraction.

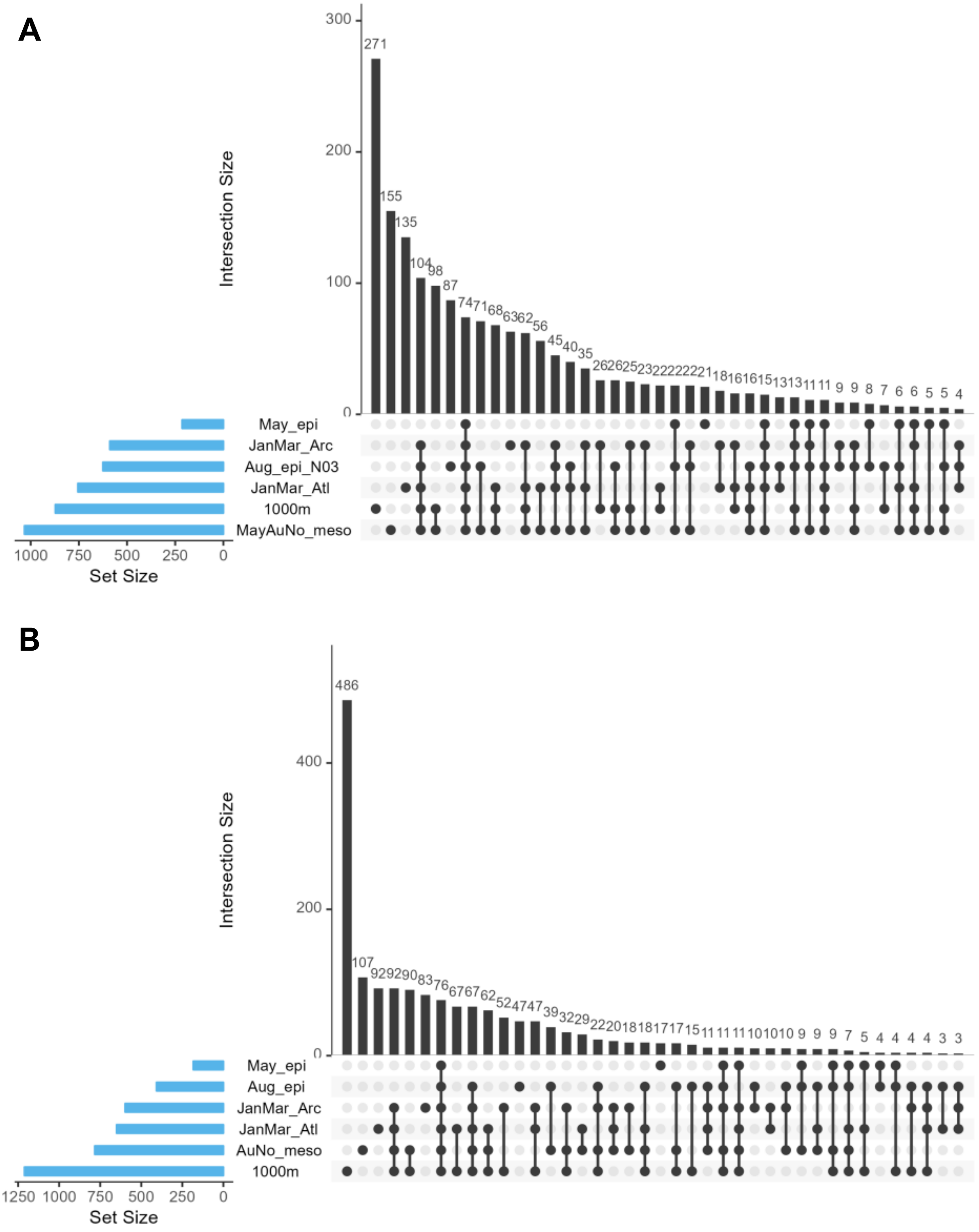


**Figure S5: UpSet plots depicting the number of shared ASVs between sample clusters** (A) Shared ASVs in the pico (0.4-3 $\mu$m) fraction, between the clusters of samples delimited in Figure **??**. (B) Shared ASVs in the nano-micro (3-200 $\mu$m) fraction.
